# First inhibitor of a bacterial two-partner secretion system

**DOI:** 10.64898/2026.01.11.698920

**Authors:** Alfred Hartojo, Laurence Don Wai Luu, Lachlan Adamson, Kyra Majors, Ashleigh S. Paparella, Peggy A. Cotter, Richard M. Johnson, Matthew Thomas Doyle

**Affiliations:** Sydney Infectious Disease Institute, The University of Sydney, Darlington, New South Wales, Australia; School of Medical Sciences, Faculty of Medicine and Health, The University of Sydney, Darlington, New South Wales, Australia; Centre for Drug Discovery Innovation, The University of Sydney, Darlington, New South Wales, Australia; School of Biotechnology and Biomolecular Sciences, University of New South Wales, Sydney, NSW, 2052, Australia; Department of Microbiology and Immunology, School of Medicine, University of North Carolina-Chapel Hill, Chapel Hill, North Carolina, USA; Department of Microbiology and Immunology, East Carolina University, Brody School of Medicine, Greenville, North Carolina, USA

## Abstract

Two-partner secretion system transporter proteins (TpsB) are widely conserved across Gram-negative pathogens. TpsB family proteins secrete exoprotein virulence factors that perform a myriad of functions such as adhesion and immune modulation. Despite their incredible importance in bacterial infectious disease, TpsB inhibitors have not yet been discovered. Here, we describe a potent inhibitor of FhaC, a TpsB protein produced by *Bordetella spp*. FhaC secretes the exoprotein FhaB that is essential for the establishment of whooping cough. We designed a peptide called P1 that we predicted would prevent substrate binding and lock FhaC in a secretion-inactive state. Simulations and biochemical assays supported our hypothesis and identified interactions important for P1 binding to FhaC. Strikingly, we observed that the peptide strongly inhibited FhaB secretion from clinical isolates and broadly reduced correlates of virulence. Together, this work provides a strong case for further development of a novel class of anti-TpsB anti-virulence compounds.

## INTRODUCTION

Despite the accelerating antibiotics resistance crisis, there is a waning pipeline of novel antibiotics in development (1, 2). Antibiotic resistant infections caused by Gram-negative species disproportionately contribute towards death statistics which is thought to be due to their high intrinsic resistance imparted by the presence of an outer membrane (OM) that acts as a barrier against antibiotic entry (3, 4). An alternative approach to developing traditional antibiotics is to develop inhibitors of virulence factors that bacteria use to cause acute and persistent infections. Unlike broad-spectrum antibiotics that disrupt healthy human microbiomes and pose a high selective pressure for the acquisition of specific resistance, anti-virulence compounds are likely to be more specific to pathogens (5). However, the high species specificity of inhibitors currently under investigation that target virulence factors such as toxins, type 3 secretion systems s, or proteins involved in quorum sensing pathways, are expected to narrow the utility of these inhibitors to a small subset of clinical disease indications (5–7).

<u>T</u>wo-<u>p</u>artner <u>s</u>ecretion protein <u>B</u> (TpsB) transporters are a novel target for the development of anti-virulence compounds with wider clinical utility against a broad spectrum of Gram-negative pathogens. TpsB transporters are amongst the most common secretion systems of those found only within Gram-negative bacteria and are conserved in all major pathogenic species relevant to human health (8, 9). TpsB proteins are embedded in the OM and function to export ‘exoprotein’ virulence factors across the OM (8). Essential virulence functions of TpsB-secreted exoproteins include attachment to biotic/abiotic surfaces, biofilm stabilization, interbacterial competition, toxin delivery, nutrient scavenging, and host-immune evasion (8, 10, 11). Significant progress has been made to understand the structure and functional mechanism of TpsB proteins (12–16).

Currently solved structures of TpsB transporters are essentially superimposable; their common architecture consists of an N-terminal α-helix (H1), a linker region, two periplasmic polypeptide transport-associated (POTRA) domains, and an OM embedded β-barrel domain into which H1 is inserted (**Figure 1A**) (13–15). Current models posit that H1 acts as a plug when the TpsB transporter is at rest but exits the β-barrel channel when a nascent exoprotein substrate binds to conserved grooves on the POTRA domains (**Figure 1**, step 1) (12, 15–17). The exoprotein can then be translocated through the channel to the cell surface, wherein conserved TpsB surface loops 3 and 4 act to accelerate folding and translocation to the extracellular space (**Figure 1**, steps 2-3) (16). Although TpsB transporters are members of the Omp85 superfamily that includes outer membrane protein insertases such as the bacterial BamA and mitochondrial Sam50 proteins, TpsB structures diverge significantly (9, 18–20). This finding suggests that TpsB inhibitors that lack Omp85 cross-reactivity can be developed.

**Figure 1:**
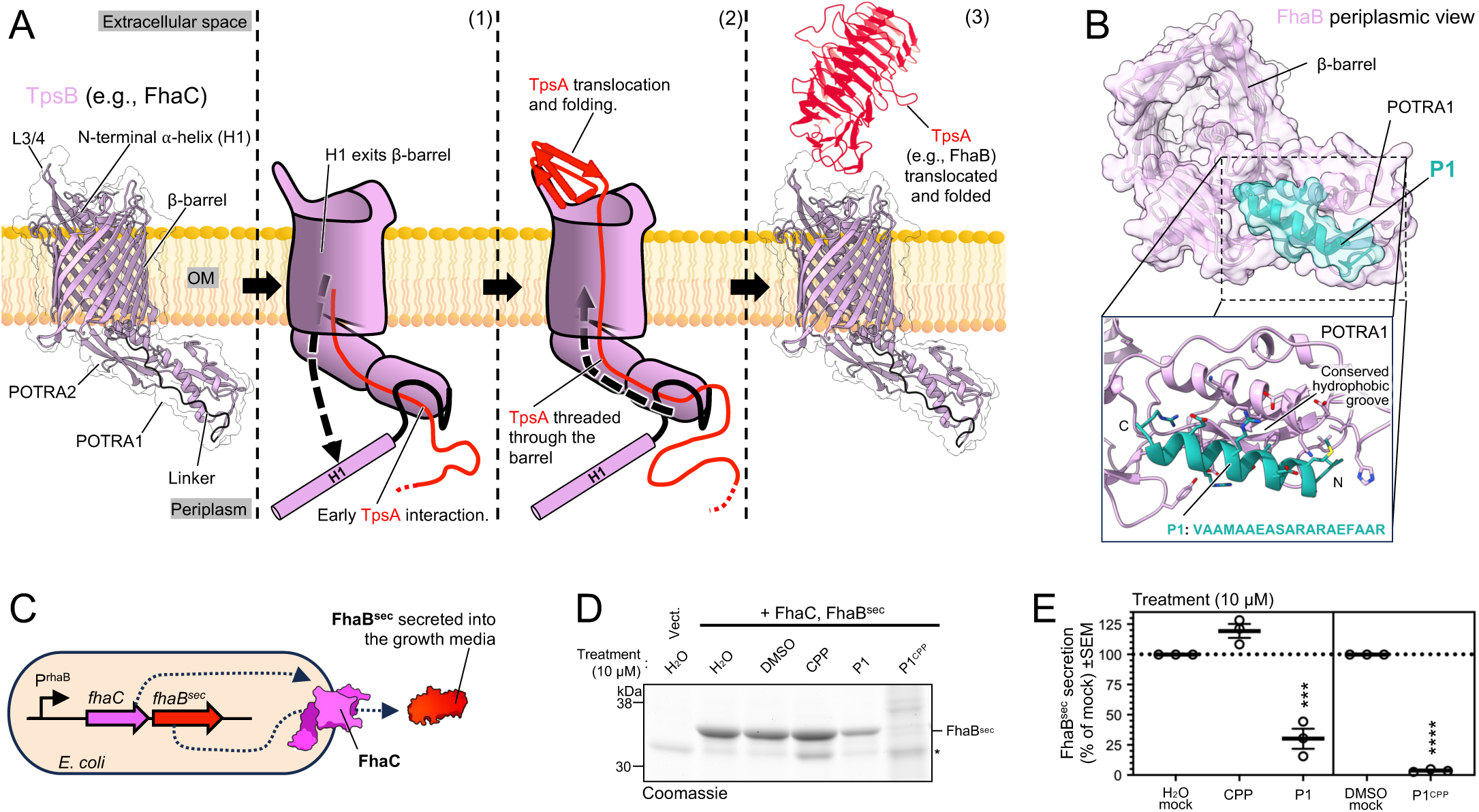
Two-partner secretion and design of an inhibitor of FhaB exoprotein secretion. (**A**) Model of exoprotein secretion by TpsB transporter family proteins. TpsB proteins such as *B. pertussis* FhaC have an N-terminal α-helix (H1) embedded inside the lumen of the β-barrel domain. Extracellular β-barrel loops 3 and 4 (L3/4) are surface exposed and POTRA domains 1 and 2 are present in the periplasm. H1 is linked to POTRA1 via the linker region. The FhaC structure depicted is based on PDB 3NJT(13) and AlphaFold was used to predict missing loops and linker. The FhaB (red) structure is PDB 1RWR(28). Step 1; engagement of a nascent unfolded exoprotein with the POTRA1/2 domains is thought to trigger disengagement of H1 from the β-barrel and initiation of exoprotein outer membrane (OM) translocation. Step 2; L3/4 are required for the surface folding and translocation of the exoprotein. Step 3, upon completion of an exoprotein secretion reaction a TpsB transporter is thought to return to the resting state. (**B**) Periplasmic view of an RFdiffusion-generated peptide called P1 (Table S1) predicted bound to the conserved hydrophobic groove of POTRA1 (see conservation in Figure S1). (**C**) Scheme for testing potential FhaC inhibitors. *E. coli* harboring a plasmid with *fhaC* and *fhaB^sec^* genes in an operon under the control of a rhamnose inducible promoter (P^RhaB^). When the TPS system is active, FhaB^sec^ is secreted into the culture media by FhaC. (**D**) Representative experiment as described in **C**. Cultures were treated with potential inhibitors or their diluents (mock treated). Proteins in culture supernatants were separated by SDS-PAGE and detected by Coomassie strain. An unknown endogenously released *E. coli* protein is noted by (*). (**E**) Three experiments conducted as in **D** and secreted FhaB^sec^ quantified as a mean percentage of mock treatment group. ANOVA statistics in Table S8.

Here, we discovered the first inhibitor of a TpsB transporter. We focused on the most well characterized TpsB family member, FhaC, which is 97% identical across *Bordetella spp.* (8, 13, 16). FhaC exports the multifunctional exoprotein FhaB that is essential for *B. pertussis* and *B. bronchiseptica* to colonize the lower respiratory tract and to evade immune responses and thereby cause persistent acute infections in humans (whooping cough) and livestock (11, 21–23). We used a generative artificial intelligence model to design potential inhibitors of FhaC which were tested for activity to yield an inhibitor, P1, that potently reduced secretion of FhaB from clinical isolates of *B. pertussis* and *B. bronchiseptica*. Computational and biochemical experiments suggested a mechanism whereby P1 treatment can stabilize FhaC in an inactive resting state through binding to the POTRA1 domain. Not only did our inhibitor block exoprotein secretion, but it also reduced the secretion of several other key *B. pertussis* virulence factors and suggests that anti-TpsB compounds have an even greater potential as anti-virulence therapeutics than we first anticipated.

## RESULTS

### Design and identification of a FhaC inhibitor

We hypothesized that TpsB transporters would be vulnerable to inhibition by small peptides designed to compete for the binding of exoprotein substrates to the transporter. To this end, we used the generative protein diffusion program RFdiffusion (24) to design peptides *de novo* to bind to known functional epitopes of FhaC (**Table S1**). This yielded four potential domain-specific high-ranking FhaC binders; (**P1**) a peptide designed to bind to the conserved groove on the periplasmic POTRA1 domain (TpsB-specific POTRA1 Pfam hidden Markov model (HMM): PF08479 (9)), (**P2**) a peptide designed to bind to the conserved groove on POTRA2 (TpsB-specific POTRA2, PF03865), and peptides (**L_3/4_v1** & **L_3/4_v2**) designed to bind to β-barrel surface loops 3 and 4 (**Figure 1B,S1 & S2A**). We reasoned that **P1** and/or **P2** might antagonize early steps in the engagement of pre-secreted FhaB exoprotein with FhaC (13, 25), whereas **L_3/4_v1** and/or **L_3/4_v2** might prevent FhaB surface folding that contributes to its efficient translocation across the OM (16). Given the location of target POTRA domains in the periplasm, we also designed derivatives of **P1** and **P2** fused to a carrier <u>c</u>ell-<u>p</u>enetrating <u>p</u>eptide (CPP) (Table S2), that was previously shown to facilitate transport of cargo peptides across bacterial OMs (26), yielding **P1^CPP^** and **P2^CPP^**.

To test the potential FhaC inhibitors we developed a whole-cell secretion assay. We transformed *E. coli* with a low-copy plasmid harboring an operon containing codon-optimized *fhaC* and a C-terminally truncated *fhaB* allele (Δ467-3590) that encodes the exoprotein (FhaB^sec^) under the control of a *rhaB* promoter (27) (**Figure 1C**). We predicted that FhaB^sec^ (∼37kDa) would be efficiently secreted by FhaCand released into the culture medium like other truncated derivatives of FhaB (28). As expected, SDS-PAGE resolved FhaB^sec^ as the major protein released by the strain into the culture medium post-induction (**Figure 1D & S2B)**. Next, we conducted an initial screen of our peptides for inhibitory activity wherein a high peptide concentration of 10 μM was added to cultures immediately prior to induction of expression of FhaC-FhaB^sec^. Strikingly, we observed that **P1** reduced FhaB^sec^ secretion into the culture medium by ∼70% relative to a mock treated cells and that **P1^CPP^**abolished secretion (**Figure 1D,E & S2C**). We did not observe any inhibitory activity associated with treatment with the control CPP peptide or the putative inhibitor peptides **P2**, **L_3/4_v1**, or **L_3/4_v2** which were therefore excluded from further analysis (**Figure S2C,D,E**). Interestingly, we also observed that **P1^CPP^** treatment at 10 μM resulted in a reduction in final culture density (**Figure S2F**). This aligns with our observation of significant release of other proteins into the culture medium upon treatment with 10 μM **P1^CPP^** (**Figure 1D & S2C**) and together suggests that at these concentrations the inhibitor can cause bacterial cell integrity defects alongside a reduction in FhaB virulence factor secretion. We did not observe a reduction in culture density or release of other proteins into the culture medium upon treatment with 10 μM **CPP** (**Figure S2C,F**) which suggests that these effects are imparted by an increase in activity associated with the P1 moiety of **P1^CPP^**.

### P1 inhibitor acts in the periplasm

The finding that **P1^CPP^** had greater inhibitory activity at 10 μM than **P1**, and that treatment of bacteria with CPP alone did not reduce FhaB^sec^ secretion, suggested that the inhibitory portion of the peptide must cross the OM to inhibit FhaC (**Figure 1D,E**). This is consistent with our design of **P1** as a binder of the periplasmic FhaC POTRA1 domain. Additionally, the sequence of **P1** is amphipathic which might provide it with intrinsic OM penetrating function and explain the partial inhibitory activity of **P1** without the CPP carrier sequence (**Figure 1B,D,E**). To investigate the mechanism of FhaC inhibition by **P1/P1^CPP^**, we first repeated the whole-cell secretion assay with multiple concentrations to determine the concentrations at which each compound reduce FhaB^sec^ secretion by 50% (IC_50_). We found that the IC_50_ of **P1** and **P1^CPP^** for FhaB^sec^ secretion inhibition was 9.22 ± 0.16 µM and 0.68 ± 0.07 µM, respectively (**Figure 2A**). At a concentration of 2 µM **P1^CPP^** almost completely abolished FhaB^sec^ secretion, whereas 2 µM **P1** treatment had no effect. These data strongly suggest that the higher potency of **P1^CPP^** is due to its ability to cross the OM barrier. Additionally, no growth defects or non-specific release of proteins into the growth media were observed in response to any of the **P1/P1^CPP^** concentrations tested in the titration series (**Figure S3**). We therefore chose to conduct subsequent mechanistic experiments using 2 µM **P1^CPP^**.

**Figure 2:**
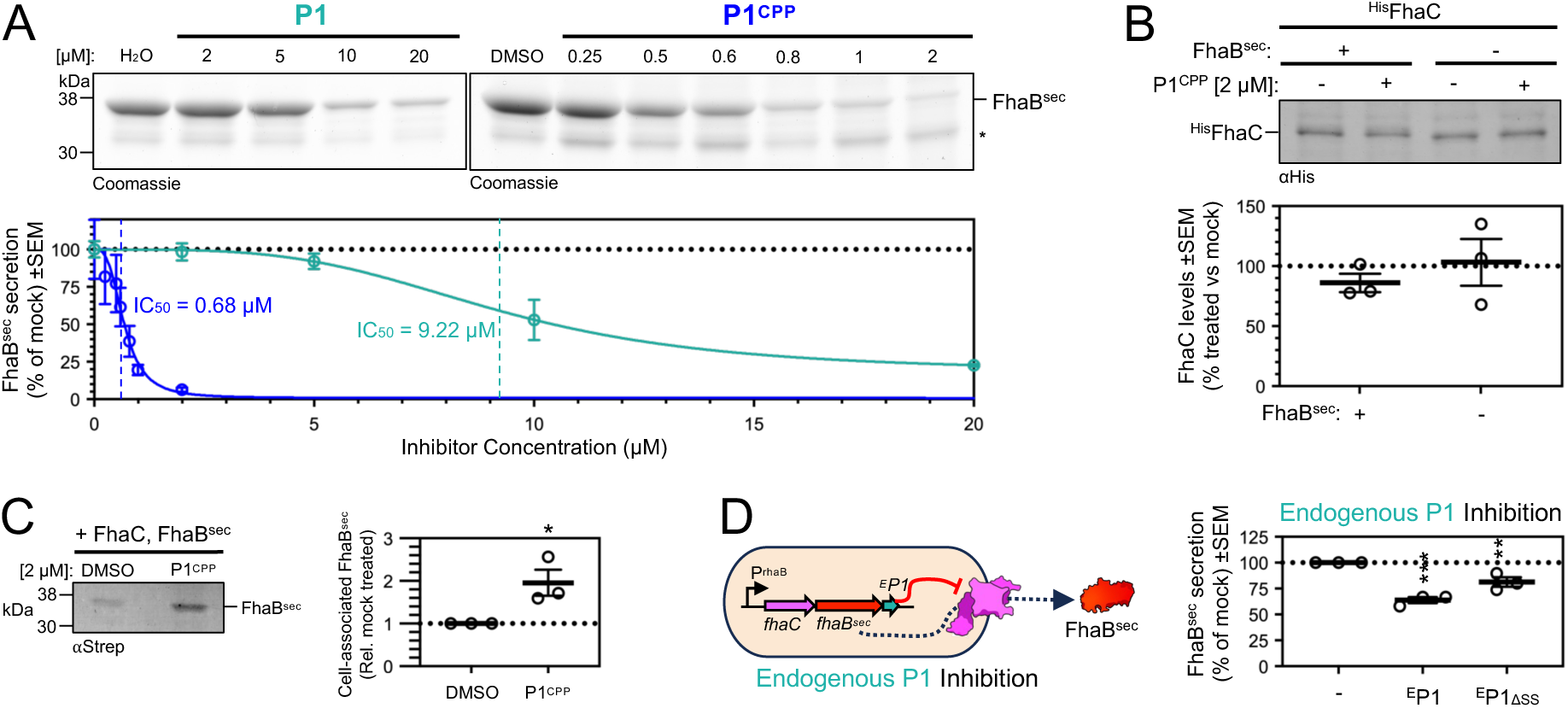
P1 inhibitor acts in the periplasm. (**A**) FhaB^sec^ secretion experiment as in Figure 1B-E except using varied concentrations of inhibitor. Top, representative gels (n = 3). An unknown endogenously released *E. coli* protein is noted by (*). Bottom, secreted FhaB^sec^ quantified as a mean percentage of mock treatment group. Fitted sigmoidal curve (R^2^ = 0.99 and 0.97, respectively) suggests 50% inhibition concentration (IC_50_) of 9.22 μM and 0.68 μM for P1 and P1^CPP^, respectively. (**B**) P1^CPP^ does not change FhaC expression. *E. coli* producing ^His^FhaC and FhaB^sec^, or just ^His^FhaC, where mock treated or treated with P1^CPP^ and total cell proteins analyzed by ⍺His western immunoblot, n = 3. (**C**) P1^CPP^ causes cellular accumulation of FhaB^sec^. Experiment conducted as in **B** except supernatant proteins analyzed by ⍺StrepII western immunoblot to detect FhaB^sec^. (**D**) Endogenously produced P1 inhibits FhaB^sec^ secretion. Experiment conducted as in Figure 1C except that a synthetic gene encoding for the P1 peptide is also included in the *fhaC* and *fhaB^sec^* operon. ΔSS = a version of the gene in which the signal sequence is removed. Student’s t-tests and ANOVA statistics for all panels in Table S8.

We next conducted several experiments that provided evidence that **P1^CPP^** inhibits the export activity of FhaC by binding to its periplasmic domains. To exclude the possibility that **P1^CPP^** treatment prevents FhaB^sec^ secretion simply by reducing expression of the FhaC transporter, we grew *E. coli* producing an N-terminally His-tagged version of FhaC (^His^FhaC) in either the absence or presence of co-expression of FhaB^sec^. These cultures were treated with **P1^CPP^**, or mock treated, and then ^His^FhaC levels were detected by western immunoblotting of total bacterial protein extracts with an αHis antibody (**Figure 2B**). As expected, regardless of FhaB^sec^ co-expression, treatment with **P1^CPP^**had no effect on the level of ^His^FhaC relative to mock-treated controls (**Figure 2B**). These data strongly suggest that **P1^CPP^** treatment blocks the function but does not alter the abundance, of the FhaC transporter. Such a scenario predicts that **P1^CPP^** inhibition of FhaC would cause a cellular accumulation of untranslocated FhaB^sec^ substrate.

Indeed, treatment of the FhaC-FhaB^sec^ expression strain with **P1^CPP^** and subsequent immunoblot of total protein extract with αStrepII (to detect a C-terminal Twin-StrepII tag on FhaB^sec^) showed accumulation of FhaB^sec^ relative to a mock treated control (**Figure 2C**). To test if periplasmic accumulation of FhaB^sec^ triggers a cell envelope stress response we monitored levels of the key periplasmic protease DegP that degrades misfolded proteins (29), the periplasmic chaperone SurA that delivers outer membrane protein substrates to the POTRA domains of BamA for OM insertion (29), and BamA. **P1^CPP^** treatment had no significant effect on steady state levels any of these proteins (**Figure S4**) suggesting that periplasmic accumulation of truncated FhaB^sec^ due to treatment with 2 μM P1^CPP^ does not induce significant stress responses in *E. coli*. As an aside, these data align well with our observation that **P1^CPP^** treatment does not affect FhaC production, including its assembly into the OM by BamA, and therefore indicates that the inhibitor is unable to cross-inhibit the divergent POTRA domains of BamA under our experimental conditions. To further test the periplasmic activity of the **P1** peptide, we also constructed a derivative of the *fhaC*-*fhaB^sec^* operon plasmid that contains a synthetic gene encoding the endogenous production of P1 (^E^P1) (**Figure 2D, left**). ^E^P1 also contains an N-terminal signal peptide enabling its secretion into the periplasm by the Sec translocon. We observed a ∼35% reduction in FhaB^sec^ secretion upon ^E^P1 co-expression (**Figure 2D, right**); a striking effect given that ^E^P1 is unlikely to achieve saturating concentrations in the periplasm relative to FhaC assuming a 1:1 production from the operon. Upon removal of the signal sequence (^E^P1_ΔSS_) FhaB^sec^ secretion was inhibited to a lesser extent (∼19%) indicating that the localization of ^E^P1 to the periplasm is important for function but that the peptide likely possesses intrinsic membrane permeation properties to allow its translocation across the inner membrane post-synthesis.

### P1 activity is dependent on the FhaC helix-linker region and conformational stabilization

Our above results are consistent with a model in which **P1** competes for initial substrate engagement to the POTRA1 groove. Previous work has suggested that substrate binding to POTRA1 can induce displacement of H1 from the transporter channel as a prerequisite for secretion (15, 17). We hypothesized that our inhibitor could stabilize the conformation of FhaC because the putative **P1**-POTRA1 binding site is located proximal to the linker region and H1. To test our hypothesis, we produced ^His^FhaC in *E. coli*, treated the bacteria with **P1^CPP^,** and probed the live bacterial cells with αHis and a fluorescent secondary antibody to detect the proportion of FhaC molecules with a surface-exposed H1 that is accessible to the antibodies (**Figure 3A**). We observed that **P1^CPP^** treatment caused a significant increase in fluorescence relative to mock-treated bacteria (**Figure 3B**). To ensure that treatment with **P1^CPP^** does not allow the antibodies to bind to periplasmic targets, we included a control experiment in which the ^His^FhaC producing *E. coli* strain was probed with antibodies specific for the periplasmic DegP protein. The α-DegP antibody labeling resulted in no detectable fluorescence supporting this approach to distinguish between surface exposed and periplasmic protein domains (**Figure S5**). The data suggest that **P1^CPP^** treatment increases the proportion of FhaC molecules in an inactive state in which the channel of the transporter is occluded by the H1 helix. This model therefore predicts that the activity of **P1^CPP^**is dependent not only on its binding to the POTRA1 groove but also its stabilization of H1, perhaps through interactions with the linker between H1 and the rest of FhaC. To test this idea, we constructed two variants of the ^His^FhaC-FhaB^sec^ (WT) expression strain that produce domain deletion variants of FhaC (**Figure 3C**); one in which H1 is deleted (ΔH1) and another that lacks both H1 and the linker (ΔH1L) and then compared FhaB^sec^ secretion from each strain in the absence or presence of **P1^CPP^** treatment. As expected, deletion of H1, or H1 and the linker, reduced the secretion of FhaB^sec^ by ∼85% (**Figure 3D**), consistent with previous reports that these domains are important for FhaC function (30, 31). Importantly, however, ^His^FhaC^ΔH1^ and ^His^FhaC^ΔH1L^ still retain some secretion function. Although **P1^CPP^**treatment reduced FhaB^sec^ secretion from strain ΔH1, the magnitude of the inhibitory effect on ΔH1 was half that of the WT strain, and there was no inhibitory effect on ΔH1L (**Figure 3D**, top and inset). Immunoblots of total bacterial protein extracts that ^His^FhaC, ^His^FhaC^ΔH1^, and ^His^FhaC^ΔH1L^ are expressed at similar levels, suggesting that the effects are due to a loss in the function of the transporter rather than its production (**Figure 3D**, bottom). By showing that **P1^CPP^** requires the presence of the linker for its activity, the data suggest that the peptide might function by forming direct interactions that stabilize H1 within the FhaC channel.

**Figure 3:**
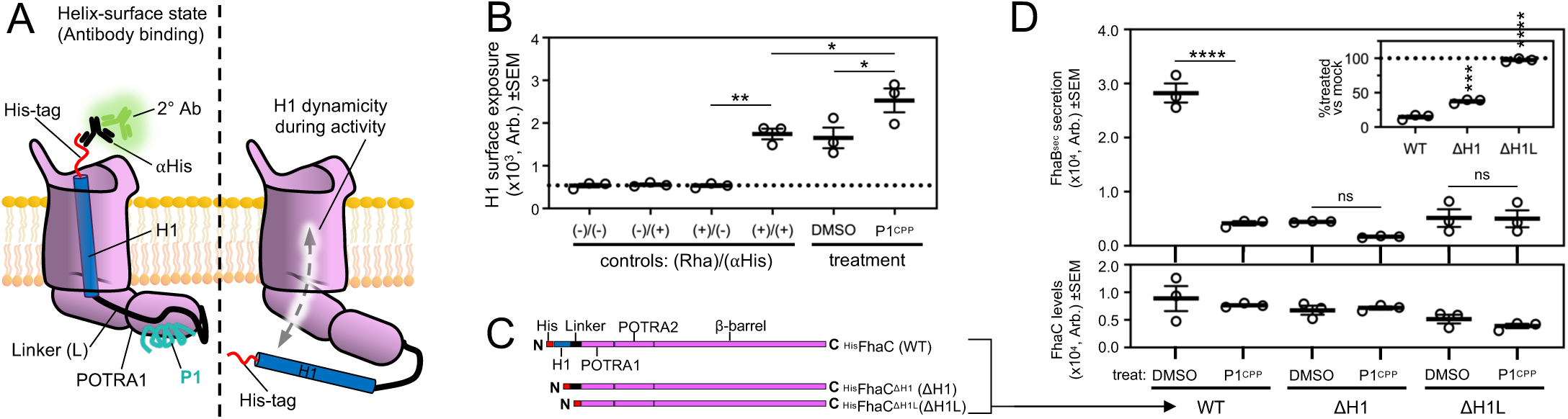
P1 activity is dependent on FhaC helix-linker region movements. Depiction of live-cell helix-mobility assay. Left, ^His^FhaC could be in a helix-surface exposed state wherein an N-terminal His-tag can be detected. Right, conformational changes in the helix/linker could prevent detection of the His tag. (**B**) P1^CPP^ treatment increased H1 surface exposure. *E. coli* producing ^His^FhaC where mock treated or treated with **P1^CPP^**(2 μM) for 30 min, and then surface exposed His-tags where monitored as in **A** (n = 3). Induction and antibody negative and positive controls were included. (**C**) Scheme of FhaC and deletion mutant primary structures. (**D**) Helix and linker deletions as in **C** reduce baseline FhaB^sec^ secretion but decrease relative potency of inhibitor. *E. coli* producing FhaB^sec^ and either ^His^FhaC (WT) or ^His^FhaC^ΔH1^ (ΔH1) or ^His^FhaC^ΔH1L^ (ΔH1L) truncated derivatives of the transporter mock treated or treated with P1^CPP^ (2 μM) as in Figure 2B, n = 3. Top, quantitation of absolute FhaB^sec^ secretion monitored by Coomassie gel of culture supernatant. Inset, FhaB^sec^ secretion upon P1^CPP^ treatment relative to mock treated. Bottom, absolute levels of ^His^FhaC and derivatives monitored by αHis immunoblot of total cellular protein. ANOVA statistics for all panels in Table S8.

### Structure-activity relationship of the FhaC inhibitor

To gain insight into the likely binding mechanism that results in FhaC inhibition by **P1**, we conducted a suite of all-atom molecular dynamics (MD) simulations. First, we used AlphaFold2 (AF2) to predict the structure of **P1** bound with that of POTRA1, both POTRA1 and POTRA2 domains together, or POTRA2. For **P1**:POTRA1, AF2 predicted **P1** bound to the conserved groove interacting with both β-sheet and α-helix residues, as expected (**Figure 4A**). Similarly, when both POTRA domains were present, AF2 preferentially predicted **P1** bound to the POTRA1 groove (**Figure 4B**). When POTRA2 was the only domain provided, AF2 predicted **P1** bound in perpendicular orientation to the strands of the β-sheet, but with an interface predicted template modelling score (iPTM) much lower than the **P1**:POTRA1 prediction (0.135 vs 0.702) (**Figure 4C**). Triplicate MD simulations on these models also suggested that the **P1**:POTRA1 binding mode might be more stable than the **P1**:POTRA2 binding mode. In these experiments we monitored the distance between a backbone α-carbon near the midpoint of P1 and a proximal α-carbon in the binding site of the respective POTRA domain based on the assumption that these distances are unlikely to vary during a stable binding interaction. The association of the measured atoms was observed to be extremely stable for **P1**:POTRA1 and **P1**:POTRA1/2 (**Figure 4A,B**) whereas the mid-point of **P1** was highly dynamic relative to POTRA2 which suggests an unstable binding interface (**Figure 4C**). Together the data indicate that **P1** can potentially bind to both POTRA domains of FhaC but that stable inhibition is likely to be dependent on binding to POTRA1.

**Figure 4:**
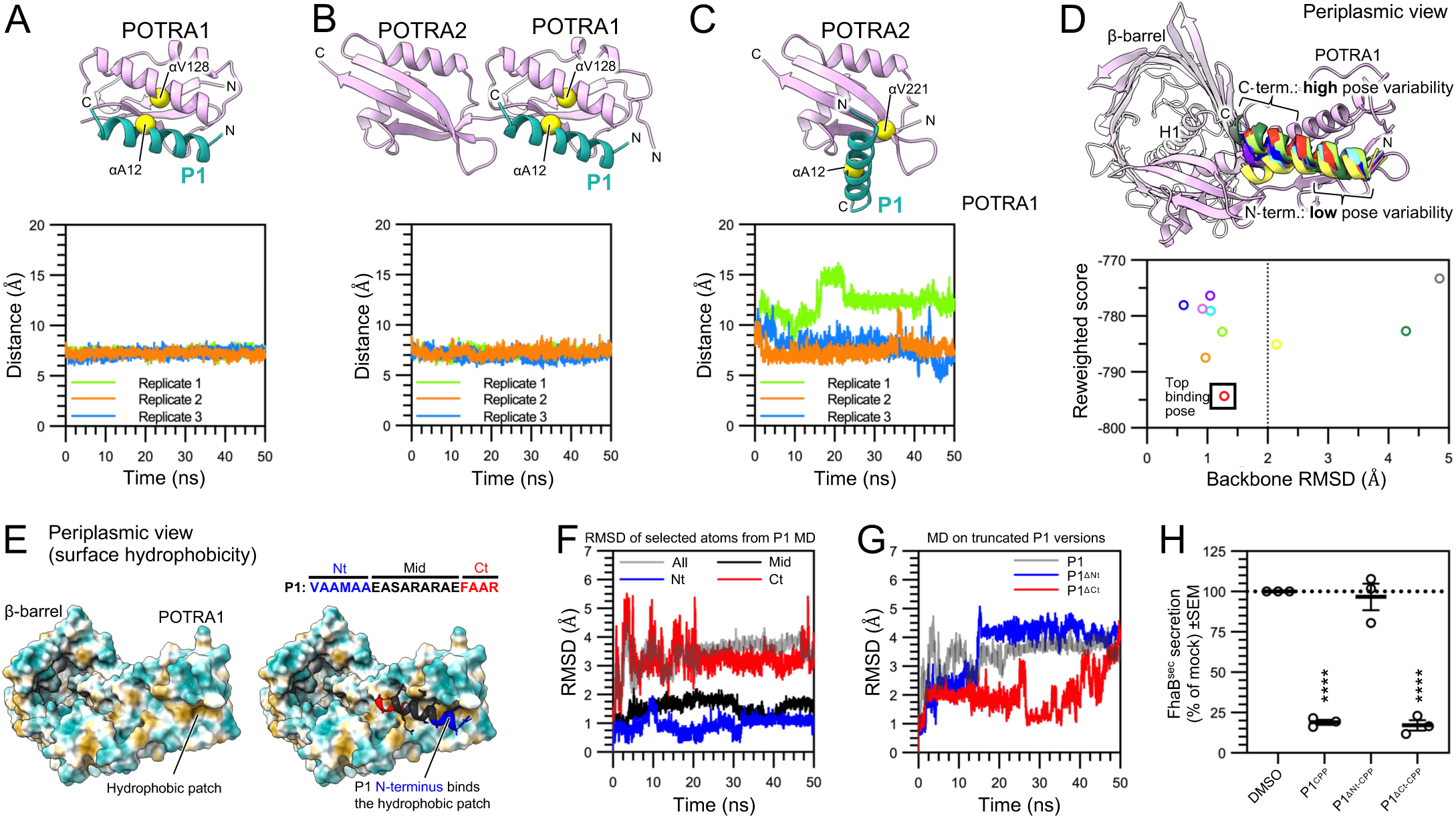
The N-terminus of P1 is required for inhibition of FhaC. (**A**) All-atom MD simulation of stable P1:POTRA1 binding. Top; AlphaFold2-predicted model of P1 bound to the POTRA1 domain of FhaC. iPTM = 0.702. Bottom; MD simulations on the predicted model. Distance between alpha carbons of P1(A12) and FhaC(V128) (yellow spheres) were monitored as a measure of biding stability. (**B**) Experiment as in **A** except using a predicted model of P1 bound to POTRA1/2. iPTM = 0.905. (**C**) Experiment as in **A** except using a predicted model of P1 bound to POTRA2 and the distance was measured between P1(A12) and FhaC(V221). iPTM = 0.135. (**D**) Consistent docking poses of the middle and N-terminus of P1 with the POTRA1 groove using FlexPepDock. Top, Structure prediction of top 10 binding poses. Bottom, energy landscape of top 10 binding poses. (**E**) Solvent-excluded protein surface of FhaC colored by hydrophilicity (teal) or hydrophobicity (tan). Left, apo; Right, top-ranked P1:POTRA1 binding pose from **D**. (**F**) Most stable binding by middle and N-terminal regions of P1. Experiment as in **A** except RMSD was monitored relative to frame 1 for all P1 atoms (grey) or only atoms in regions of P1 colored in **E**. (**G**) Deletion of C-terminal P1 residues does not affect inhibitor stability. (**H**) N-terminal P1 residues are necessary for inhibitory activity. Experiment conducted as in Figure 3D, except bacteria were treated with treated with **P1^CPP^**, its truncated derivatives (**P1^ΔNt^-^CPP^**or **P1^ΔCt^-^CPP^**), or mock-treated. ANOVA statistics in Table S8.

Further analysis revealed the importance of N-terminal residues in **P1** for its inhibitory activity. We used Rosetta FlexPepDock (32) to identify variance in binding energetics for different potential binding poses in the **P1**:POTRA1 interface (**Figure 4D**). We found that the C-terminus of **P1** varied significantly across potential binding poses and that the best scoring poses most closely aligned with a hydrophobic patch in the conserved groove of POTRA1 (**Figure 4D,E**). Indeed, when we monitored changes in **P1** conformational variation via root mean square deviation (RMSD) we noticed that although the N-terminus remained conformationally stable, significant conformational instability was observed in the C-terminus of the peptide (**Figure 4F**). Simulations in which we monitored stability of variants of **P1** containing N- or C-terminal deletions (residues indicated in **Figure 4E**) also indicated that the C-terminal residues play a minimal role in stabilizing P1 and suggests that this portion of the inhibitor might not contribute to its activity (**Figure 4G**). To test this idea, we tested **P1^CPP^** variants **P1^ΔNt-CPP^** and **P1^ΔCt-CPP^** in our FhaB^sec^ secretion assay. Consistent with our simulations, we observed that the deletion of N-terminal residues resulted in a complete loss of activity (**Figure 4H**, P1^ΔNt-CPP^) whereas deletion of C-terminal residues had no effect (**Figure 4H**, P1^ΔCt-CPP^). These data suggest that the N-terminus of **P1** is necessary for stable binding to FhaC and that the C-terminal residues are dispensable.

Our MD simulations also suggested that three potential electrostatic interactions formed between the middle section of **P1** and POTRA1 might contribute to binding stability (**Figure 5A**). Monitoring the distance between the potential salt bridges P1(R11):FhaC(E122) (**Figure 5A**, pink), P1(R13):FhaC(E162) (**Figure 5A**, purple), and P1(R15):FhaC(K129) (**Figure 5A**, brown) suggested that the interactions between R11:E122 and R13:E162 might be more stable and important for activity than E15:K129 (**Figure 5B**). To test our hypothesis, we tested charge-reversed variants of **P1^CPP^** in our FhaB^sec^ secretion assay. Consistent with our hypothesis, **P1^R11E-CPP^** and **P1^R13E-CPP^** possessed markedly lower inhibitory activity than **P1^CPP^** (**Figure 5C**). Interestingly, **P1^E15R-CPP^** did not have reduced activity but rather had a slightly greater potency than **P1^CPP^** (IC_50_ 0.56 ± 0.01 µM vs 0.68 ± 0.07 µM, respectively) (**Figure 5C**). The negative effects on inhibitory activity for R11E and R13E were consistent across double (**Figure 5D**) and triple (**Figure 5E**) substitution variants; the loss of function from the R11E substitution was particularly dominant over the positive effect of E15R. Further analyses of the simulations suggest that **P1**(R11) is part of a triad of residues that interchangeably interact with each other (**Figure S6)**. **P1**(R11) is predicted to form an alternative salt bridge interaction with another **P1** residue (E7) and a hydrogen bond with FhaC(N121), while **P1**(E7) also forms a hydrogen bond with FhaC(N121) (**Figure S6**), which likely explains why the R11E substitution results in an almost complete loss of inhibition.

**Figure 5:**
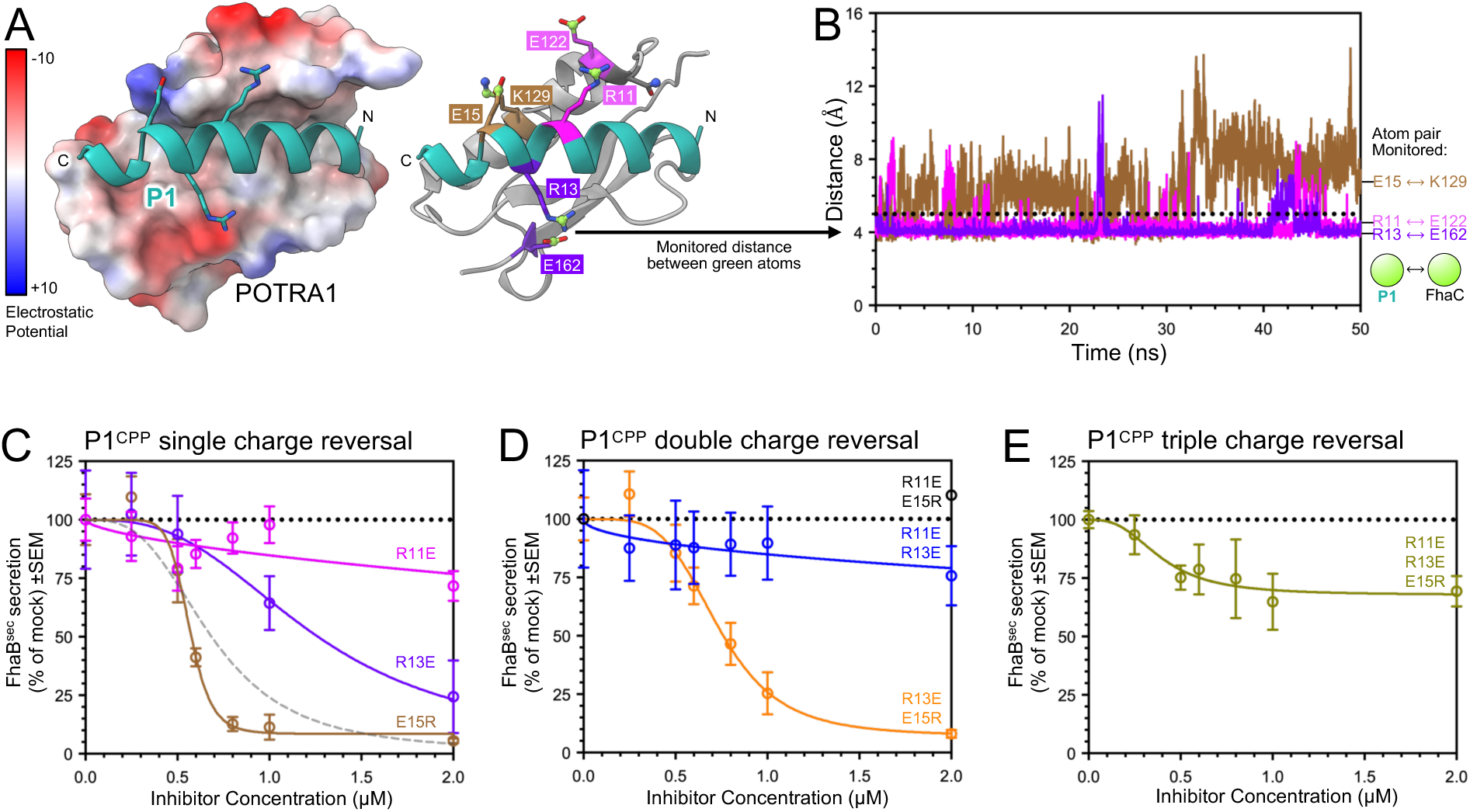
Electrostatic interactions contribute to P1 activity. (**A**) Left, solvent-excluded protein surface of POTRA1 colored by electrostatic potential. Right, predicted residue electrostatic interactions between P1 and POTRA1 colored by pair. (**B**) Experiment as in Figure 4A except distance between terminal residue carbons of residue pairs as depicted as green spheres in **A**. Dotted line denotes the maximum distance to form a stable salt bridge interaction. (**C**) FhaB^sec^ secretion experiment as in Figure 2A except using varied concentrations of single charge reversal variants of P1^CPP^. Gray dashed line indicates curve from P1^CPP^ data, Figure 2A. IC_50_ were calculated for datasets that fitted a sigmoidal curve with R^2^ > 0.95. R13E, R^2^= 0.99, IC_50_ = 1.23 µM; E15R, R^2^= 0.99, IC_50_ = 0.56 µM. (**D**) Experiment as in **C** except double charge reversal variants of P1^CPP^ were investigated. R13E/E15R, R^2^= 0.98, IC_50_ = 0.74 µM. (**E**) Experiment as in **C** except a triple charge reversal variant of P1^CPP^ was investigated. ANOVA statistics in Table S8.

### P1 broadly disrupts *Bordetella* correlates of virulence

We investigated the activity of our FhaC inhibitor against several relevant isolates of *Bordetella spp*.; *B. bronchiseptica* RB50 which is a veterinary isolate and widely studied reference strain (33), and *B. pertussis* strains L1423, L1756, and L2228 which are clinical isolates from the Australian 2008-2012 and 2013-2017 epidemics (34, 35). The *B. pertussis* strains were chosen based on their allelic variation in secreted virulence factors. L1423 [*prn*^+^/*fhaB*^+^] produces both FhaB/C and the adhesin pertactin(*prn*), L1756 [*prn*^-^/*fhaB*^+^] produces only FhaB/C, and L2228 possesses the rarely isolated *prn*^-^/*fhaB*^-^ genotype that produce neither virulence factor (35).

**P1^CPP^** not only inhibited the release of natively produced and processed FhaB (FHA) from all tested strains but also reduced the release of several other key virulence factors. First, we treated *B. bronchiseptica* RB50 with **P1^CPP^** in broth culture and observed a marked reduction in FHA secretion when we probed the growth media by immunoblot (**Figure 6A**, left). We also probed these samples with an antiserum raised against adenylate cyclase toxin (ACT), another key virulence factor that was recently identified to be loaded onto FhaB as a toxin-delivery mechanism for defense against phagocytic cells (11). Strikingly, we did not observe any ACT in the growth medium upon **P1^CPP^**treatment (**Figure 6A**, right). We similarly treated the *B. pertussis* isolates and likewise observed significant reductions in FHA secretion and a corresponding accumulation of the exoprotein in whole cell lysate (**Figure 6B**). We conducted untargeted protein identification by LC-MS/MS on L1423 culture supernatants after mock or **P1^CPP^** treatment. Remarkably, alongside a reduction in FhaB secretion due to **P1^CPP^** treatment as expected, we also observed a reduction in the release of several major virulence factors including the autotransporter Vag8 which functions in serum resistance (36, 37) and subunits PtxA and PtxD of the pertussis toxin complex that is critically involved in immune suppressive functions and leukocytosis (38, 39) (**Figure 6C**, red; **Table S7**). We observed an increase in the levels of several proteins released into the media including SspA and ArgD which are cytoplasmic proteins involved in stress responses and arginine synthesis (**Figure 6C**, green**; Table S7**). Additionally, we observed 137 cytoplasmic and periplasmic proteins that were only detected in the growth media upon **P1^CPP^** treatment that are largely involved in metabolic processes (**Table S7**). Together with our observation that high concentrations of the inhibitor cause significant release of *E. coli* proteins into the growth media, these data indicate that treatment of *B. pertussis* with **P1^CPP^** results in cell envelope stress. We also analyzed protein levels in **P1^CPP^**treated L1423 whole cell pellets and found no significant differences relative to untreated bacteria (**Table S7**).

**Figure 6:**
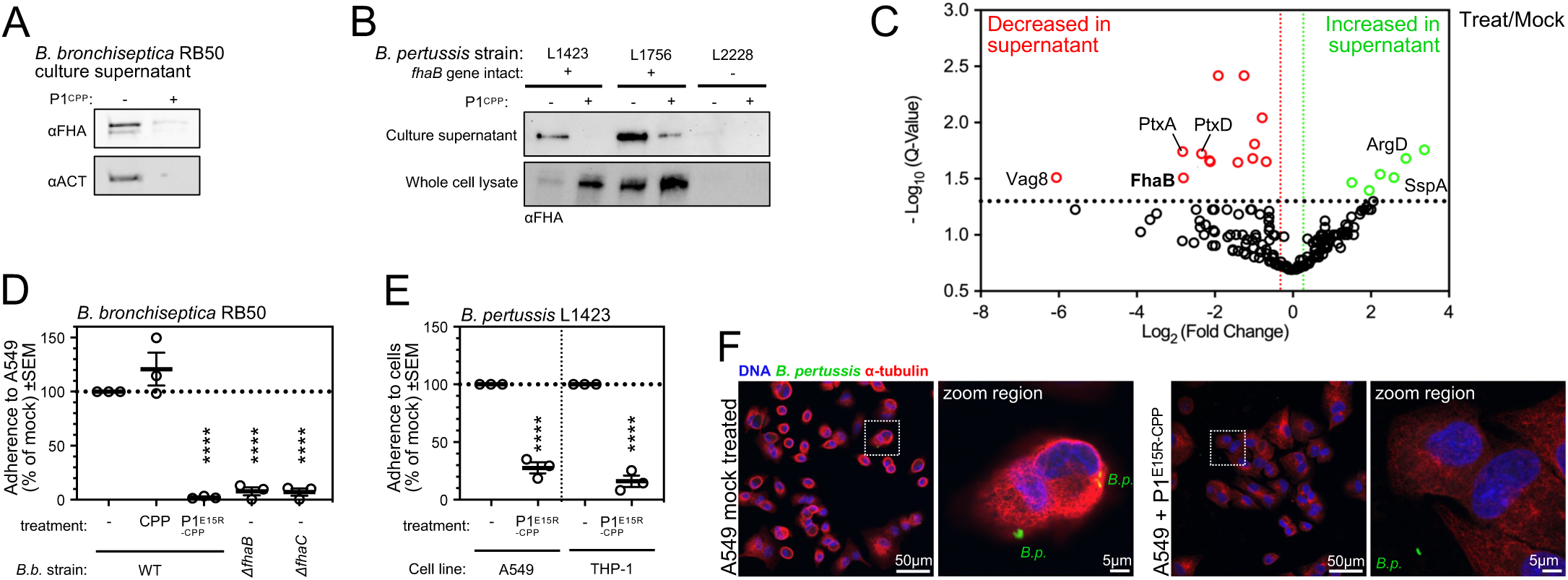
P1 inhibits Bordetella bronchiseptica and Bordetella pertussis virulence functions. (**A**) P1^CPP^ prevented secretion of FHA and ACT from *B. bronchiseptica* RB50. RB50 was grown in non-virulence factor repressive conditions before transfer to virulence inducing conditions and treatment with P1^CPP^ (10 μM) for 1h (see *Methods*), n = 3. Proteins released to culture media were probed by immunoblotting with either ⍺FHA or ⍺ACT antisera. (**B**) P1^CPP^ inhibited secretion of FHA from *B. pertussis* clinical isolates. Strains L1423, L1756, and L2228 were grown similarly to **A** except bacteria were treated with P1^CPP^ (20 μM) for 2h. Proteins released to culture media and whole cell lysates were probed by immunoblotting with ⍺FHA. (**C**) P1^CPP^ treatment reduces virulence factor release from *B. pertussis*. Volcano plot of LC-MS/MS data showing differential protein levels in the supernatant (n = 3). Red symbols indicate proteins of significantly lower abundance in the culture media of P1^CPP^ treated cells. Green symbols indicate proteins of significantly higher abundance. Vertical lines indicate fold-change cutoff >20%. Horizontal line indicates q-value cutoff at 0.05. Full proteomics datasets are in Table S7 (**D**) P1^E15R^-^CPP^ prevents *B. bronchiseptica* adherence to lung cells. Adherent RB50 infection of A549 cell line in the presence of inhibitor treatment (or mock treatment) quantified by recovered colony count, n = 3. Isogenic RB50 *ΔfhaB* and RB50 *ΔfhaC* strains were included as controls. (**E**) P1^E15R^-^CPP^ prevents *B. pertussis* adherence to lung cells and macrophages. Adherent L1423 after infection of either A549 lung or macrophage differentiated THP-1 cell lines in the presence of inhibitor treatment (or mock treatment) quantified by microscopy, n = 3. L1423 bacteria were transformed with a plasmid expressing fuGFP. (**F**) P1^E15R^-^CPP^ prevents *B. pertussis* induced cytotoxicity and adherence. Representative micrographs from infected A549 cells in **E** are shown. DNA was labelled with DAPI, and α-tubulin was labelled with a corresponding antiserum. Sample micrographs of treated infected THP-1 cells are in Figure S8. ANOVA statistics for all panels in Table S8.

Finally, we observed that **P1^CPP^** significantly reduced the ability of *Bordetella spp*. to bind to host cells and protected them from bacterial induced cytotoxicity. In these experiments the A549 lung epithelial cell line was infected with bacteria simultaneously with treatment with the more potent **P1^E15R-CPP^** derivative of the inhibitor. By measuring recovered cfu we observed a 97% loss in relative adherence of *B. bronchiseptica* RB50 to A549 cells upon **P1^E15R^-^CPP^** treatment (**Figure 6D**). This effect phenocopied the loss of adherence observed for isogenic knock out strains RB50*ΔfhaB* and RB50*ΔfhaC* (**Figure 6D**). As expected, treatment with **^CPP^** had no effect on RB50 adherence to A549 cells (**Figure 6D**). By microscopic analysis, we also observed that **P1^E15R-CPP^** treatment resulted in a ∼75% loss in adherence of *B. pertussis* L1423 to A549 cells, and an ∼85% loss in adherence to macrophage-differentiated THP-1 cells (**Figure 6E**). Mock-treated A549 and THP-1 cells infected with *B. pertussis* possessed a rounded morphology and a peripheral reordering of tubulin indicating that L1423 infection was cytotoxic in the absence of FhaC inhibitor treatment (**Figure 6F & S7**). The cytotoxicity phenotype was not observed for **P1^E15R-CPP^**-treated cells (**Figure 6F & S7**). These data not only suggest that **P1^E15R^-^CPP^** treatment can prevent bacterial-induced cytotoxicity but also that the inhibitor itself is non-toxic to A549 and THP-1 cells under our experimental conditions.

## DISCUSSION

37 years since the discovery of the widespread two-partner secretion system (40), this report describes the discovery of the first inhibitor of a TpsB transporter. The recent breakthrough in the success rates for computational design of protein-target binders encouraged us to use the AI software RFdiffusion (24, 41, 42) to design short peptides to bind to conserved and functionally important features of FhaC, a TpsB transporter essential for the virulence of *Bordetella* spp (43). **P1**, a peptide designed to bind to the conserved groove of the FhaC POTRA1 domain, showed remarkable inhibitory activity against the secretion of FhaB from *B. pertussis*, *B. bronchisceptica* and the recombinant derivative FhaB^sec^ from *E. coli*. All-atom simulations suggested that **P1** binds stably to the POTRA1 groove via both hydrophobic and electrostatic interactions and subsequent experiments using **P1** derivatives designed to disrupt these interactions consistently yielded reduced activity. However, peptides that we designed to bind to the conserved groove in POTRA2, or surface loops 3 and 4 of the β-barrel domain, had no detectible activity. It remains possible that all of these alternative peptides failed to bind to FhaC altogether. Alternatively, the peptides might bind weakly without causing a defect in FhaC function and, perhaps in the case of the putative loop binders, are easily displaced by the emerging FhaB exoprotein as it folds on the surface loops by β-augmentation (16). We therefore do not exclude the possibility that conserved TpsB features other than the POTRA1 groove are viable targets for inhibitor design that might require alternate inhibitor scaffolds or the development of surface binding antibodies (44).

Our results suggest that the **P1** peptide can inhibit multiple aspects of the FhaC secretion cycle. Given previous work that showed that mutations in the POTRA1/2 domain grooves reduced the ability of FhaC to bind to substrates in vitro (12) and our observation that FhaB accumulated inside cells upon **P1** treatment, **P1**:POTRA1-groove binding might therefore compete with the binding of nascent periplasmic exoprotein substrates to the transporter. Our model is consistent with (1) our observation that treatment with **P1** increased the level of surface exposed FhaC H1 α-helix, (2) previous crystal structures of TpsB proteins solved in secretion inactive states wherein H1 is surface exposed and occludes the β-barrel channel (13–15), and (3) PELDOR spectroscopy experiments that show that substrate binding is required for removal of H1 from the channel (17). We also observed that the efficacy of **P1** was diminished when H1, or both H1 and the linker, were deleted, and that our FlexPepDock analysis predicted interactions between the N-terminus of **P1** and the FhaC linker. It is therefore plausible that **P1**:POTRA1 binding further stabilizes the inactive state of FhaC by increasing the stability of the linker similarly to the previously isolated “FhaC_DIS_” mutants whose secretion activity is abolished due to increased stabilization of linker-POTRA interactions (12, 15).

Our inhibitor broadly disrupted virulence activities of both *B. bronchiseptica* and *B. pertussis* clinical isolates. This suggests that the CPP carrier peptide was not only able to carry the active **P1** moiety across the short chain lipooligosaccharide layer of the *B. pertussis* OM, but that it was also able to cross the long chain lipopolysaccharides of *B. bronchiseptica* which is otherwise thought to provide resistance against antimicrobial peptides (45, 46). We observed a reduction in FhaB secretion from both species and a concomitant loss of adherence to lung and macrophage cell lines, consistent with the adherence function of FhaB (47, 48). However, our unexpected observation that the inhibitor reduced the secretion of toxins (*B. bronchiseptica* ACT and *B. pertussis* subunits PtxA and PtxB) and serum resistance factor Vag8, strongly suggest that TpsB inhibitors have a greater than expected clinical potential to broadly inhibit virulence. It is likely that the reduction in secretion of additional virulence factors is a secondary effect. In this model, FhaB that is not secreted accumulates in the periplasm and cause a disruption in cell envelope maintenance pathways which likely impact other virulence factor secretion systems. This model is consistent with our observation that treatment with the FhaC inhibitor caused release of cytoplasmic and periplasmic proteins into culture media presumably due to resultant bacterial cell envelope leakiness. Vag8 reduction by FhaC inhibition might cause the loss of *Bordetella* serum resistance and reduce the ability of these species to persist in the respiratory tract during protracted infections. Regardless, our data strongly suggest that FhaC inhibition, and TpsB inhibition more generally, is an even more exciting approach than previously anticipated. Given the high conservation of the TpsB POTRA1 domain, in subsequent studies it will be interesting to test whether **P1** can broadly inhibit TpsB transporters across bacterial species as a path toward a novel class of pan-anti-virulence therapeutics.

## MATERIALS AND METHODS

### Design of FhaC-binding peptides

RFdiffusion (24) was used to design peptide binders of FhaC. AF2-predicted structures and crystal structures of FhaC (PDB: 3NJT and 4QKY) were used as targets in RFdiffusion to design 10-30 residue-long peptides to bind to indicated FhaC hotspots (**Table S1**). Peptides with most favorable scoring criteria (pLDDT ≥ 0.5, pAE ≤ 10, and RMSD ≤ 4 Å) were then synthesized by Genscript at >90% purity (**Table S2**).

### Bacterial strains, cell lines, and growth conditions

All strains are listed in **Table S3**. *E. coli* strains were grown overnight from a single colony at 37 ℃ with shaking at 250 rpm in lysogeny broth (LB, Lennox). *B. bronchiseptica* and *B. pertussis* strains were grown overnight from a single colony at 37 ℃ with shaking at 250 rpm in Stainer-Scholte (SS) media containing 1% heptakis (49). Antibiotics were added as appropriate; 50 µgmL^-1^ trimethoprim, 100 µgmL^-1^ ampicillin, or 50 µgmL^-1^ kanamycin. A549 cells were cultured in Dulbecco’s Modified Eagle Medium (DMEM) media, or Ham’s F12 media, with 10% fetal bovine serum (FBS), 1% penicillin-streptomycin (1000 U/ml) and 4 mM L-glutamine. THP-1 cells were cultured in Roswell Park Memorial Institute (RPMI) 1640 media with 10% FBS, penicillin-streptomycin (1000 U/ml), 1% sodium pyruvate, and 0.3% sodium bicarbonate. All cells were incubated at 37°C with 5% CO_2_.

### Plasmid construction

Plasmids used in this study are listed in **Tables S4** and oligonucleotides and gene fragments listed in **Tables S5** and **S6**, respectively. To construct pRJ64, an *E. coli* codon-optimized FhaC encoding region with a N-terminal 10xHis-tag inserted between the signal sequence and the start of H1 was cloned into the EcoRI and BamHI sites in pTrc99a. To construct pMTDS109, an *E. coli* codon-optimized operon encoding FhaC and a C-terminally truncated FhaB (Δ467-3590) that encodes the exoprotein (FhaB^sec^) also containing an N-terminal TwinStrepII tag was cloned into NdeI and XbaI sites in pSCRhaB2. To construct pMTDS263, pMTDS109 was digested with restriction enzymes HindIII and XbaI an ligated to similarly digested ^E^P1-encoding fragment mtd185. To construct pMTDS630, pRJ64 was digested with restriction enzymes NcoI and XbaI and the ^His^FhaC-encoding fragment was ligated into similarly digested pSCRhaB2. To construct pMTDS323, a 10x His-tag was introduced into pMTDS109 with primers mtd191/192. Plasmid expressing signal sequence deletion mutant of pMTDS263 (pMTDS608) was constructed via Gibson Assembly after amplification with primers mtd363/372 and mtd364/371. For deletion mutants of pMTDS323, pMTDS497 (ΔHL) was constructed by amplification with primers mtd227/228 and pMTDS607 (ΔH) by amplification with primers mtd363/370 and mtd364/369 and subsequent reassembly.

### Construction of B. bronchiseptica RB50 ΔfhaC

RB50 *ΔfhaC* was constructed by allelic exchange mutagenesis as previous (50). Briefly, to construct pRJ94 *fhaB* was amplified using overlap PCR using primers RJ481-485, and the resulting locus with an engineered deletion of aa18-576 encoding region was cloned into pSS4245 allelic exchange plasmid via Gibson Assembly. pRJ94, was introduced into RB50 by conjugation, and cointegrants were selected on BG agar containing kanamycin, streptomycin, and 50 mM MgSO_4_. Mutagenesis was confirmed by Sanger sequencing.

### *E. coli* FhaB^sec^ secretion assay

Overnight cultures of *E. coli* BL21(DE3) containing pMTDS109 (or other relevant plasmids) were subcultured into LB beginning at an OD_600_ of 0.05 and grown for 2h (37 ℃, 250rpm). The bacteria were then simultaneously supplemented with appropriate concentrations of treatments and 0.2% L-rhamnose (w/v) to induce expression of FhaB^sec^ and FhaC. The bacteria were then grown for another hour (37 ℃, 250rpm). An OD_600_ reading was taken, and the remaining bacterial cultures were centrifuged (3,220 x g, 10 min, 25 °C). The supernatant was filter-sterilized (0.22 µm) and then proteins from 5 mL aliquots of the supernatant were precipitated by the addition of 10% (v/v) trichloroacetic acid (TCA) and 4 mM phenylmethanesulfonyl fluoride (PMSF) and incubation on ice for 1h. Precipitates were then pelleted (20,817 x g, 10 min, 4 °C), washed twice with 600 μL of acetone, and air-dried (20 min, 37 ℃). 40 µL of 2x SDS Protein Loading Solution (100 mM Tris-HCl pH 8, 20% glycerol (v/v), 0.2% bromophenol blue (w/v), 4% SDS (v/v)) was added to precipitates and heated to 99 ℃ for 15min with shaking at 1200rpm. Protein samples (2 μL) were separated by SDS-PAGE on 8%–16% Tris-glycine gradient gels (ThermoFisher) at room temperature for 2 h at 150 V, stained by Coomassie Brilliant Blue (R-250), and imaged with an Odyssey DLx infrared imager (Licor) using maximum quality and resolution settings and a 700 nm laser. Intensity values of a band corresponding to FhaB^sec^ were quantified using FIJI ImageJ (v2.14.0/1.54f) and normalized to mock-treated controls.

### Whole bacterial protein samples

Strains were grown as in the *E. coli* FhaBsec secretion assay except that after induction 1 mL culture aliquots were TCA-precipitated as above. Precipitates were then resuspended in 2x SDS Protein Loading Solution in a volume normalized to an OD_600_ measurement recorded immediately as subculture samples were taken (volume in μL = 200 x OD_600_). Samples (5 µL) were then probed by Western Immunoblotting.

### Antibodies and Antisera

Monoclonal mouse anti-His, mouse anti-StrepII, and mouse anti-alpha-tubulin were obtained from Genscript (cat#A00186), Qiagen (cat#34850), and Sigma-Aldrich (cat#T5168) respectively. Goat anti-mouse 800CW IRDye (cat#926-32210) and goat anti-rabbit Ig 680LT IRDye (cat#926-680210) was obtained from Li-Cor, and goat anti-mouse IgG (H+L) – Alexa Fluor 568 was obtained from ThermoFisher (cat#A-11004). Anti-FHA, anti-ACT (35, 51, 52). Rabbit anti-DegP raised against DegP(S201A), anti-SurA raised against His-tagged SurA, and anti-BamA_C_ raised against a C-terminal BamA peptide are previously described (53, 54)

### Western immunoblotting

Western blot was conducted as previously (55). Briefly, after proteins were separated by SDS-PAGE (as above) they were transferred to nitrocellulose membranes using an iBlotII transfer device (Life Technologies). Membranes were blocked overnight with immunoblotting buffer [1:1 Intercept Blocking Buffer (Li-Cor):PBS + 0.01% Tween-20], incubated overnight with 1° antibody diluted in immunoblotting buffer (αHis or αStrep at 1:5,000 dilution; αBamA, αDegP, or αSurA at 1:10,000 dilution), washed with PBS-T (PBS + 0.01% Tween-20), incubated with 2° antibody (1:5,000 dilution) for 2 h, and finally washed two times with PBS-T and three times with PBS. Membranes were air-dried and then imaged with an Odyssey infrared as above with 700 nm and 800 nm lasers. Band intensities were quantified in FIJI as above.

### Helix mobility assay

Subcultures were grown exactly as in the *E. coli* FhaB^sec^ secretion assay except bacteria were induced 0.2% L-rhamnose (w/v) and treated with 2 μM **P1^CPP^** or mock treated for 30 mins. After treatment, the OD_600_ was recorded and 100 µL aliquots of the culture were incubated with anti-His (dilution factor of 1:100) for 30 min at 37 °C. The bacteria were then pelleted (2,000 x g, 2 min, 25 °C), supernatants removed, and bacterial pellet resuspended in 100 µL of PBS containing goat anti-mouse 800CW IRDye (dilution factor of 1:100). Bacteria were then incubated for a further 30 minutes at 37 °C before being pelleted (2,000 x g, 2 min, 25 °C), supernatants removed, washed with 1 mL PBS, and re-pelleted (2,000 x g, 2 min, 25 °C). Bacterial pellets were finally resuspended with 100 µL PBS and loaded onto 96-well, flat, clear-bottom black polystyrene microplates (Corning, cat#3603). A Varioskan Lux microplate reader (ThermoFisher) was used to measure endpoint OD_600_ values and then the microplate was scanned with maximum quality and resolution settings with an Odyssey infrared scanner at 800nm. Well intensity values were quantified in FIJI and normalized to their respective endpoint OD_600_ value.

### All-atom molecular dynamics (MD) simulations

We used AF2 to obtain structural predictions of **P1** (and its derivatives) bound and unbound to FhaC POTRA domains that were subsequently amber relaxed (56). Simulations were performed using GROMACS (57) and AMBER99SB-ILDN force field (58). Structures were solvated with the TIP3P water model (59) and 150 mM NaCl, and energy minimized using the steepest descent algorithm, the “Verlet” scheme for neighbor searching for 50,000 steps (maximum step size of 0.01), and a tolerance of 1000. A 10 Å cut-off was used for Coulombic interactions and a 10 Å cut-off for short-range Van der Waals interactions. Electrostatics were calculated using the Fast Smooth Particle-Mesh Ewald method (60). Simulations were NVT equilibrated with starting velocities assigned and position restraints, followed by NPT equilibration with position restraints before subsequently releasing the position restraints in another NPT equilibration step. A set of parameters was used for all equilibration and production steps. The leap-frog algorithm was used for motion integration, all bonds with H-atoms were constrained, the Verlet scheme was used for neighbor searching, the temperature was set at 300 K (with a modified Berendsen thermostat), and pressure was set at 1 bar (using Parrinello-Rahman pressure coupling), number of groups set for thermocoupling was two (protein and non-protein), step length was set at 2 fs. Each equilibration step was conducted for 100 ps (50,000 steps), while the production step was conducted over 50 ns (25,000,000 steps). Data points were recorded every 10 ps (5,000 steps). UCSF Chimera (v1.17.3) was used to extract RMSD values (relative to 1^st^ frame) and distances between selected atoms from the simulations.

### B. bronchiseptica secretion assay

Overnight cultures of *B. bronchiseptica* strain RB50 were washed twice in Dulbecco’s PBS (D-PBS), resuspended in SS media, and subcultured to an OD_600_ of 1 before treatment with **P1^CPP^** peptides at a concentration of 10 µM or mock treatment with equivalent volume of DMSO. Subcultures were then grown for another hour (37°C, 200 rpm). 2 mL culture aliquots where pelleted (10,000 x g, 4 min, 4°C), culture media supernatant retained and filter-sterilised (0.22 µM filter), and finally TCA precipitated (0.4 mM PMSF, 10% TCA (v/v)) on ice for 2 h. Precipitated supernatant proteins were then washed and resuspended for Western immunoblot analysis as described above.

### B. bronchiseptica adherence assay

A549 human epithelial cells were passaged (< 12 times) in Ham’s F12 media with 10% FBS and seeded at 8 × 10^4^ cells/well into 12-well trays. *B. bronchiseptica* strains were cultured overnight at 37 °C in SS media supplemented with 50 mM MgSO_4_. Bacteria were washed to remove MgSO_4_ and resuspended in SS media supplemented with 20 μM **P1^E15R^-^CPP^** (or with equivalent volume of DMSO) before infecting A549 cells at an MOI of 100 for 45 min (37 °C, 5% CO_2_). A549 cells were then washed with 37 ℃ D-PBS (3x) to remove non-adherent bacteria. A549 cells were then lysed by incubation with 250 µL 0.05% Triton X-100 for 15 min. D-PBS was then added to lysates to a total volume of 1 mL. The A549 cell lysates were serially diluted in D-PBS and plated onto Bordet-Gengou Blood agar plates and incubated (37 °C, 72 h). Recovered cfu/sample was counted for each lysate as a measure of adherence compared to mock-treated conditions.

### *B. pertussis* adherence assay

*B. pertussis* L1423 transformed with a pBBR1MCS2-fuGFP to constitutively express free-use green fluorescent protein (fuGFP) was grown on Bordet-Gengou ( BD Scientific) agar for 3 days at 37°C. A loopful of L1423-fuGFP colonies were then resuspended in SS media supplemented with 1% heptakis and grown for 24 h shaking at 180 rpm at 37°C. Twenty-four hours prior to infection, 2 × 10^5^ A549 cells and 5 × 10^5^ THP-1 cells were seeded onto 24-well plates containing round coverslips (13 mm). THP-1 cells were induced overnight into macrophage-like cells with 5 µl of 10 µM phorbol 12-myristate-13-acetate. After overnight incubation, wells were washed with 1 ml PBS and replaced with 1 ml antibiotic-free media. Each cell line was then infected with L1423-fuGFP at an MOI of 100 and in the presence of 20 μM **P1^E15R^-^CPP^** or DMSO mock control. The plates were then centrifuged at 500 x g for 5 minutes and then incubated at 37°C with 5% CO_2_ for 2 hours. The media was then removed and each well was washed three times with PBS to remove non-adherent cells. Cells were then fixed with 1 ml of 4% paraformaldehyde for 20 min, permeabilized with 0.5% Triton X-100 for 15 min, and then blocked with 1% BSA for 1 h at room temperature. Host cells were then stained with DAPI (1:40,000) and anti-alpha-tubulin (1:3000) with goat anti-mouse IgG (H+L) – Alexa Fluor 568 secondary antibody (1:600) (1 h incubations). Coverslips were then mounted with ProLong diamond mounting solution. Images were then acquired on a Leica Stellaris 5 microscope. Five field of views were randomly selected per sample (3888 x 3888 pixels). All images were imaged at 40X objective (water immersion, NA 1.1). The resulting images were imported into ImageJ (v1.54) and the ratio of attached pertussis cells to host cells quantified.

### Proteomics

*B. pertussis* L1423 was grown and treated with 20 μM **P1^CPP^,** or DMSO mock control, as above. 10 µg of whole cell lysate protein or supernatant protein were prepared and trypsin digested as previously described. Digested peptides in 0.1% formic acid were loaded onto a QExactive Orbitrap mass spectrometer which was connected to an UltiMate 3000 high-performance nano liquid chromatography system (Thermo Fisher Scientific). The resulting raw data files were loaded into Mascot (v2.51) (Matrix Science) and searched against a custom *B. pertussis* database for protein identification (61). Protein quantification using normalized total spectral counts was then performed using Scaffold (v5.3.4). Only proteins with a minimum of two peptides, protein probability threshold > 95%, and a peptide probability threshold >99% were retained. Statistical analysis was performed using a two tailed Student’s t-test with false discovery correction performed using the Storey-Tibshirani method as implemented in the Q-value R package (62). Differential abundant proteins were defined as proteins with >20 % fold-change and Q-value < 0.05. The mass spectrometry data files have been deposited in PRIDE with data set identifier PXD070712.

## Supporting information

Supplemental Figures Tables and References

Table S7 - Proteomic Identifications

Table S8 - Statistical Analyses

## Acknowledgments

AH is supported by the University of Sydney International Stipend Scholarship and the University of Sydney International Tuition Fee Scholarship. This work was funded by a National Institute of Allergy and Infectious Diseases NIH R21 Grant (R21AI180112) to PAC, MTD, and RMJ, seed funding from the University of Sydney Infectious Diseases Institute, and seed funding from the University of Sydney Centre for Drug Discovery Innovation. MTD is supported by Australian Research Council Discovery Project (DP260100880). LDWL is supported by a National Health and Medical Research Council Emerging Leadership (EL1) Fellowship (GNT2033643) and a University of New South Wales Scientia Fellowship. A patent application has been filed covering the inhibitors described in this work (PCT/AU2025/050953). We thank Harris D. Bernstein (National Institutes of Diabetes and Digestive and Kidney Diseases) for kindly providing antisera used in this work and for thoughtful comments on the manuscript.

## Author contributions

Conceptualization: AH, MTD, PAC, RMJ

Methodology: AH, AP, LDWL, LA, MTD, PAC, RMJ

Investigation: AH, KM, LA, LDWL, RMJ

Visualization: AH, MTD, RMJ

Supervision: AP, MTD, PAC, RMJ

Writing-original draft: AH

Writing-review & editing: AH, AP, LDWL, LA, MTD, PAC, RMJ

## Competing interests

All authors declare they have no competing interests.

## Notes

### Competing Interest Statement

The authors have declared no competing interest.

## REFERENCES

1. Butler MS, Vollmer W, Goodall ECA, Capon RJ, Henderson IR, Blaskovich MAT. A Review of Antibacterial Candidates with New Modes of Action. ACS Infectious Diseases. 2024;10(10):3440–74.

2. Naghavi M, Vollset SE, Ikuta KS, Swetschinski LR, Gray AP, Wool EE, et al. Global burden of bacterial antimicrobial resistance 1990-2021: a systematic analysis with forecasts to 2050. The Lancet. 2024;404(10459):1199–226.

3. Murray CJL, Ikuta KS, Sharara F, Swetschinski L, Robles Aguilar G, Gray A, et al. Global burden of bacterial antimicrobial resistance in 2019: a systematic analysis. The Lancet. 2022;399(10325):629–55.

4. Maher C, Hassan Karl A. The Gram-negative permeability barrier: tipping the balance of the in and the out. mBio. 2023;14(6):e01205–23.

5. Lau WYV, Taylor PK, Brinkman FSL, Lee AHY. Pathogen-associated gene discovery workflows for novel antivirulence therapeutic development. eBioMedicine. 2023;88.

6. Ali SO, Yu XQ, Robbie GJ, Wu Y, Shoemaker K, Yu L, et al. Phase 1 study of MEDI3902, an investigational anti-Pseudomonas aeruginosa PcrV and Psl bispecific human monoclonal antibody, in healthy adults. Clinical Microbiology and Infection. 2019;25(5):629.e1-.e6.

7. D’Angelo F, Baldelli V, Halliday N, Pantalone P, Polticelli F, Fiscarelli E, et al. Identification of FDA-Approved Drugs as Antivirulence Agents Targeting the pqs Quorum-Sensing System of Pseudomonas aeruginosa. Antimicrobial Agents and Chemotherapy. 2018;62(11):10.1128/aac.01296-18.

8. Guérin J, Bigot S, Schneider R, Buchanan SK, Jacob-Dubuisson F. Two-Partner Secretion: Combining Efficiency and Simplicity in the Secretion of Large Proteins for Bacteria-Host and Bacteria-Bacteria Interactions. Frontiers in Cellular and Infection Microbiology. 2017;7.

9. Heinz E, Lithgow T. A comprehensive analysis of the Omp85/TpsB protein superfamily structural diversity, taxonomic occurrence, and evolution. Frontiers in Microbiology. 2014;5.

10. Ruhe ZC, Subramanian P, Song K, Nguyen JY, Stevens TA, Low DA, et al. Programmed Secretion Arrest and Receptor-Triggered Toxin Export during Antibacterial Contact-Dependent Growth Inhibition. Cell. 2018;175(4):921–33.e14.

11. Nash ZM, Inatsuka CS, Cotter PA, Johnson RM. Bordetella filamentous hemagglutinin and adenylate cyclase toxin interactions on the bacterial surface are consistent with FhaB-mediated delivery of ACT to phagocytic cells. mBio. 2024:e0063224.

12. Delattre A-S, Saint N, Clantin B, Willery E, Lippens G, Locht C, et al. Substrate recognition by the POTRA domains of TpsB transporter FhaC. Molecular Microbiology. 2011;81(1):99–112.

13. Clantin B, Delattre A-S, Rucktooa P, Saint N, Méli AC, Locht C, et al. Structure of the Membrane Protein FhaC: A Member of the Omp85-TpsB Transporter Superfamily. Science. 2007;317(5840):957–61.

14. Guerin J, Botos I, Zhang Z, Lundquist K, Gumbart JC, Buchanan SK. Structural insight into toxin secretion by contact-dependent growth inhibition transporters. eLife. 2020;9:e58100.

15. Maier T, Clantin B, Gruss F, Dewitte F, Delattre A-S, Jacob-Dubuisson F, et al. Conserved Omp85 lid-lock structure and substrate recognition in FhaC. Nature Communications. 2015;6(1):7452.

16. Baud C, Guérin J, Petit E, Lesne E, Dupré E, Locht C, et al. Translocation path of a substrate protein through its Omp85 transporter. Nature Communications. 2014;5(1):5271.

17. Guérin J, Baud C, Touati N, Saint N, Willery E, Locht C, et al. Conformational dynamics of protein transporter FhaC: large-scale motions of plug helix. Molecular Microbiology. 2014;92(6):1164–76.

18. Doyle MT, Bernstein HD. Function of the Omp85 Superfamily of Outer Membrane Protein Assembly Factors and Polypeptide Transporters. Annual Review of Microbiology. 2022;76(1):259–79.

19. Diederichs KA, Ni X, Rollauer SE, Botos I, Tan X, King MS, et al. Structural insight into mitochondrial β-barrel outer membrane protein biogenesis. Nature Communications. 2020;11(1):3290.

20. Noinaj N, Kuszak AJ, Gumbart JC, Lukacik P, Chang H, Easley NC, et al. Structural insight into the biogenesis of β-barrel membrane proteins. Nature. 2013;501(7467):385–90.

21. Nash ZM, Cotter PA. Bordetella Filamentous Hemagglutinin, a Model for the Two-Partner Secretion Pathway. Microbiol Spectr. 2019;7(2).

22. Nicholson TL, Brockmeier SL, Loving CL. Contribution of Bordetella bronchiseptica filamentous hemagglutinin and pertactin to respiratory disease in swine. Infection and immunity. 2009;77(5):2136–46.

23. Melvin Jeffrey A, Scheller Erich V, Noël Christopher R, Cotter Peggy A. New Insight into Filamentous Hemagglutinin Secretion Reveals a Role for Full-Length FhaB in Bordetella Virulence. mBio. 2015;6(4):10.1128/mbio.01189-15.

24. Watson JL, Juergens D, Bennett NR, Trippe BL, Yim J, Eisenach HE, et al. De novo design of protein structure and function with RFdiffusion. Nature. 2023.

25. Hodak H, Clantin B, Willery E, Villeret V, Locht C, Jacob-Dubuisson F. Secretion signal of the filamentous haemagglutinin, a model two-partner secretion substrate. Molecular Microbiology. 2006;61(2):368–82.

26. Yamamoto K, Yamamoto N, Ayukawa S, Yasutake Y, Ishiya K, Nakashima N. Scaffold size-dependent effect on the enhanced uptake of antibiotics and other compounds by Escherichia coli and Pseudomonas aeruginosa. Scientific Reports. 2022;12(1):5609.

27. Cardona ST, Valvano MA. An expression vector containing a rhamnose-inducible promoter provides tightly regulated gene expression in Burkholderia cenocepacia. Plasmid. 2005;54(3):219–28.

28. Clantin B, Hodak H, Willery E, Locht C, Jacob-Dubuisson F, Villeret V. The crystal structure of filamentous hemagglutinin secretion domain and its implications for the two-partner secretion pathway. Proceedings of the National Academy of Sciences. 2004;101(16):6194–9.

29. Sklar JG, Wu T, Kahne D, Silhavy TJ. Defining the roles of the periplasmic chaperones SurA, Skp, and DegP in Escherichia coli. Genes & development. 2007;21(19):2473–84.

30. Guédin S, Willery E, Tommassen J, Fort E, Drobecq H, Locht C, et al. Novel Topological Features of FhaC, the Outer Membrane Transporter Involved in the Secretion of the Bordetella pertussis Filamentous Hemagglutinin*. Journal of Biological Chemistry. 2000;275(39):30202–10.

31. Guérin J, Saint N, Baud C, Meli AC, Etienne E, Locht C, et al. Dynamic interplay of membrane-proximal POTRA domain and conserved loop L6 in Omp85 transporter FhaC. Molecular Microbiology. 2015;98(3):490–501.

32. Raveh B, London N, Zimmerman L, Schueler-Furman O. Rosetta FlexPepDock ab-initio: Simultaneous Folding, Docking and Refinement of Peptides onto Their Receptors. PLOS ONE. 2011;6(4):e18934.

33. Cotter PA, Miller JF. BvgAS-mediated signal transduction: analysis of phase-locked regulatory mutants of Bordetella bronchiseptica in a rabbit model. Infect Immun. 1994;62(8):3381–90.

34. Octavia S, Sintchenko V, Gilbert GL, Lawrence A, Keil AD, Hogg G, et al. Newly Emerging Clones of Bordetella pertussis Carrying prn2 and ptxP3 Alleles Implicated in Australian Pertussis Epidemic in 2008–2010. The Journal of Infectious Diseases. 2012;205(8):1220–4.

35. Xu Z, Octavia S, Luu LDW, Payne M, Timms V, Tay CY, et al. Pertactin-Negative and Filamentous Hemagglutinin-Negative Bordetella pertussis, Australia, 2013–2017. Emerging Infectious Disease journal. 2019;25(6):1196.

36. Hovingh ES, de Maat S, Cloherty APM, Johnson S, Pinelli E, Maas C, et al. Virulence Associated Gene 8 of Bordetella pertussis Enhances Contact System Activity by Inhibiting the Regulatory Function of Complement Regulator C1 Inhibitor. Frontiers in Immunology. 2018;Volume 9 - 2018.

37. Marr N, Shah NR, Lee R, Kim EJ, Fernandez RC. Bordetella pertussis Autotransporter Vag8 Binds Human C1 Esterase Inhibitor and Confers Serum Resistance. PLOS ONE. 2011;6(6):e20585.

38. Black WJ, Munoz JJ, Peacock MG, Schad PA, Cowell JL, Burchall JJ, et al. ADP-Ribosyltransferase Activity of Pertussis Toxin and Immunomodulation by Bordetella pertussis. Science. 1988;240(4852):656–9.

39. Connelly Carey E, Sun Y, Carbonetti Nicholas H. Pertussis Toxin Exacerbates and Prolongs Airway Inflammatory Responses during Bordetella pertussis Infection. Infection and Immunity. 2012;80(12):4317–32.

40. Schiebel E, Schwarz H, Braun V. Subcellular location and unique secretion of the hemolysin of Serratia marcescens. J Biol Chem. 1989;264(27):16311–20.

41. Fox DR, Asadollahi K, Samuels I, Spicer BA, Kropp A, Lupton CJ, et al. Inhibiting heme piracy by pathogenic Escherichia coli using de novo-designed proteins. Nature Communications. 2025;16(1):6066.

42. Ragotte RJ, Liang H, Tam J, Miletic S, Berman JM, Palou R, et al. De novo design of potent inhibitors of clostridial family toxins. Proceedings of the National Academy of Sciences. 2025;122(39):e2509329122.

43. Willems RJL, Geuijen C, van der Heide HGJ, Renauld G, Berlin P, van den Akker WMR, et al. Mutational analysis of the Bordetella pertussis fim/fha gene cluster: identification of a gene with sequence similarities to haemolysin accessory genes involved in export of FHA. Molecular Microbiology. 1994;11(2):337–47.

44. Bennett NR, Watson JL, Ragotte RJ, Borst AJ, See DL, Weidle C, et al. Atomically accurate de novo design of antibodies with RFdiffusion. bioRxiv. 2025:2024.03.14.585103.

45. Di Fabio JL, Caroff M, Karibian D, Richards JC, Perry MB. Characterization of the common antigenic lipopolysaccharide O-chains produced by Bordetella bronchiseptica and Bordetella parapertussis. FEMS Microbiology Letters. 1992;97(3):275–81.

46. Banemann A, Deppisch H, Gross R. The lipopolysaccharide of Bordetella bronchiseptica acts as a protective shield against antimicrobial peptides. Infect Immun. 1998;66(12):5607–12.

47. Locht C, Berlin P, Menozzi FD, Renauld† G. The filamentous haemagglutinin, a multifaceted adhesin produced by virulent Bordetella spp. Molecular Microbiology. 1993;9(4):653–60.

48. Serra DO, Conover MS, Arnal L, Sloan GP, Rodriguez ME, Yantorno OM, et al. FHA-Mediated Cell-Substrate and Cell-Cell Adhesions Are Critical for Bordetella pertussis Biofilm Formation on Abiotic Surfaces and in the Mouse Nose and the Trachea. PLOS ONE. 2011;6(12):e28811.

49. Hulbert RR, Cotter PA. Laboratory Maintenance of Bordetella pertussis. Current Protocols in Microbiology. 2009;15(1):4B.1.-4B.1.9.

50. Inatsuka Carol S, Xu Q, Vujkovic-Cvijin I, Wong S, Stibitz S, Miller Jeff F, et al. Pertactin Is Required for Bordetella Species To Resist Neutrophil-Mediated Clearance. Infection and Immunity. 2010;78(7):2901–9.

51. Julio SM, Inatsuka CS, Mazar J, Dieterich C, Relman DA, Cotter PA. Natural-host animal models indicate functional interchangeability between the filamentous haemagglutinins of Bordetella pertussis and Bordetella bronchiseptica and reveal a role for the mature C-terminal domain, but not the RGD motif, during infection. Molecular microbiology. 2009;71(6):1574–90.

52. Lee SJ, Gray MC, Guo L, Sebo P, Hewlett EL. Epitope Mapping of Monoclonal Antibodies againstBordetella pertussis Adenylate Cyclase Toxin. Infection and Immunity. 1999;67(5):2090–5.

53. Wang X, Peterson JH, Bernstein HD. Bacterial Outer Membrane Proteins Are Targeted to the Bam Complex by Two Parallel Mechanisms. mBio. 2021;12(3):10.1128/mbio.00597-21.

54. Pavlova O, Peterson JH, Ieva R, Bernstein HD. Mechanistic link between β barrel assembly and the initiation of autotransporter secretion. Proceedings of the National Academy of Sciences. 2013;110(10):E938–E47.

55. Hanson SE, Dowdy T, Larion M, Doyle MT, Bernstein HD. The patatin-like protein PlpD forms structurally dynamic homodimers in the Pseudomonas aeruginosa outer membrane. Nature Communications. 2024;15(1):4389.

56. Evans R, O’Neill M, Pritzel A, Antropova N, Senior A, Green T, et al. Protein complex prediction with AlphaFold-Multimer. bioRxiv. 2022:2021.10.04.463034.

57. Abraham MJ, Murtola T, Schulz R, Páll S, Smith JC, Hess B, et al. GROMACS: High performance molecular simulations through multi-level parallelism from laptops to supercomputers. SoftwareX. 2015;1-2:19–25.

58. Lindorff-Larsen K, Piana S, Palmo K, Maragakis P, Klepeis JL, Dror RO, et al. Improved side-chain torsion potentials for the Amber ff99SB protein force field. Proteins: Structure, Function, and Bioinformatics. 2010;78(8):1950–8.

59. Price DJ, Brooks CL, III. A modified TIP3P water potential for simulation with Ewald summation. The Journal of Chemical Physics. 2004;121(20):10096–103.

60. Essmann U, Perera L, Berkowitz ML, Darden T, Lee H, Pedersen LG. A smooth particle mesh Ewald method. The Journal of Chemical Physics. 1995;103(19):8577–93.

61. Luu LDW, Octavia S, Zhong L, Raftery MJ, Sintchenko V, Lan R. Proteomic Adaptation of Australian Epidemic Bordetella pertussis. PROTEOMICS. 2018;18(8):1700237.

62. John D. Storey AJB, Alan Dabney, David Robinson, Gregory Warnes. qvalue: Q-value estimation for false discovery rate control. 2.43.0 ed2025.

