## Supplemental Figures Tables and References for "First inhibitor of a bacterial two-partner secretion system"

**This PDF file includes:**

Figure S1: P1 is predicted to bind the conserved hydrophobic pocket in FhaC POTRA1. (pg 2)

Figure S2: Activity of RFdiffusion-predicted peptide binders of FhaC. (pg 3-4)

Figure S3: Concentration-dependent inhibition of FhaC – full gels & $OD_{600}$ per concentration. (pg 5)

Figure S4: P1$^{CPP}$ does not alter expression of SurA, DegP, or BamA. (pg 6)

Figure S5: Live-cell helix-mobility assay specificity control. (pg 7)

Figure S6: P1 residue R11 interacts interchangeably with FhaC N121. (pg 8)

Figure S7: P1$^{E15R\text{-}CPP}$ prevents *B. pertussis* induced cytotoxicity and adherence to differentiated macrophages. (pg 9)

Table S1. RFdiffusion output scores and predicted properties of the top 10 peptide sequences of each run conducted. (pg 10-11)

Table S2. Molecular properties of P1 and derivatives. (pg 12)

Table S3. Strains used in this study. (pg 13)

Table S4. Plasmids used in this study. (pg 14)

Table S5. Single-stranded oligonucleotides used in this study. (pg 15)

Table S6. Double-stranded oligonucleotide fragments used in this study. (pg 16)

Supplementary References (pg 17)

**Other Supplementary Materials for this manuscript include the following:**

Proteomics Data Analysis (Table S7)

Statistical Significance Data (Table S8)

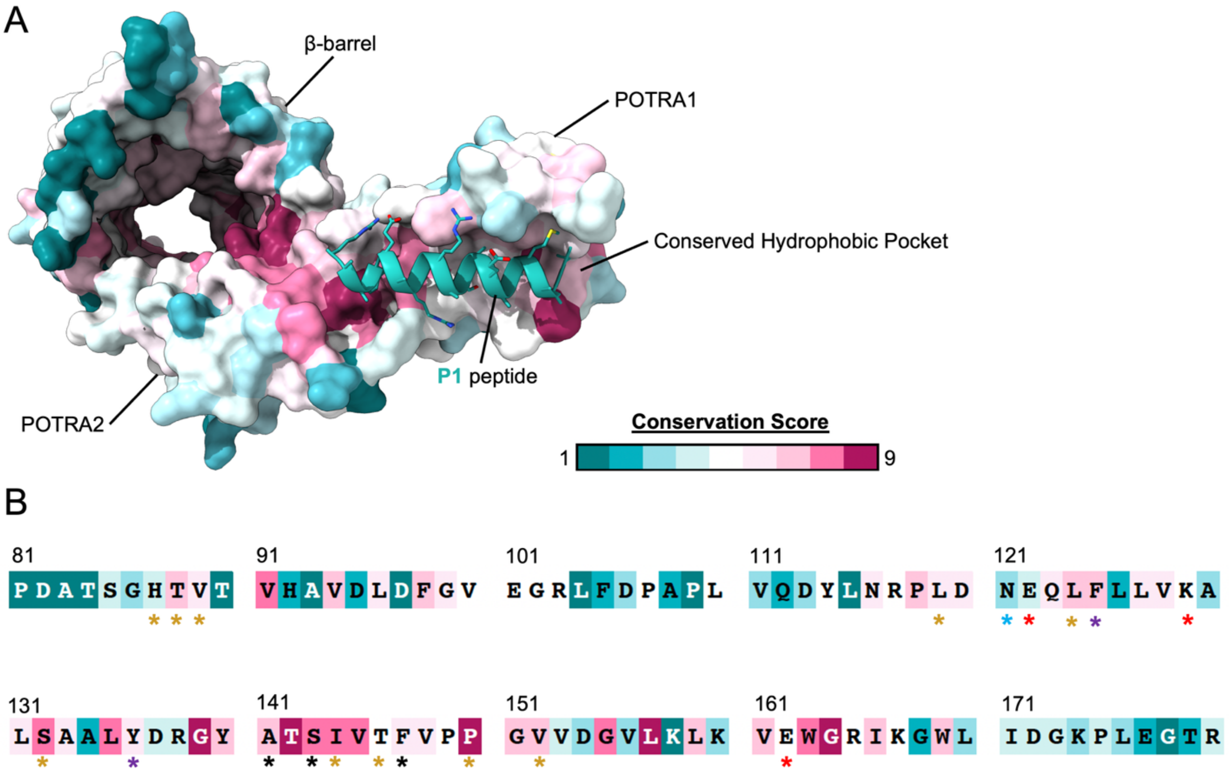


**Figure S1: P1 is predicted to bind the conserved hydrophobic pocket in FhaC POTRA1.** (**A**) Per-residue conservation of FhaC (PDB: 4QKY) (*1*) based on ConSurf Web Server scores (*2*) with P1 docked into the POTRA1 pocket. Colour depicts the conservation level of each residue across homologues of FhaC. (**B**) Sequence of FhaC POTRA1 (86-164) colored by conservation level per residue. Asterisks are colored by the type of predicted interaction with **P1**: red = salt bridge interaction, brown = contact or Van der Waals interactions, blue = hydrogen bonds, purple = cation-π interactions, black = hydrogen backbone interactions.

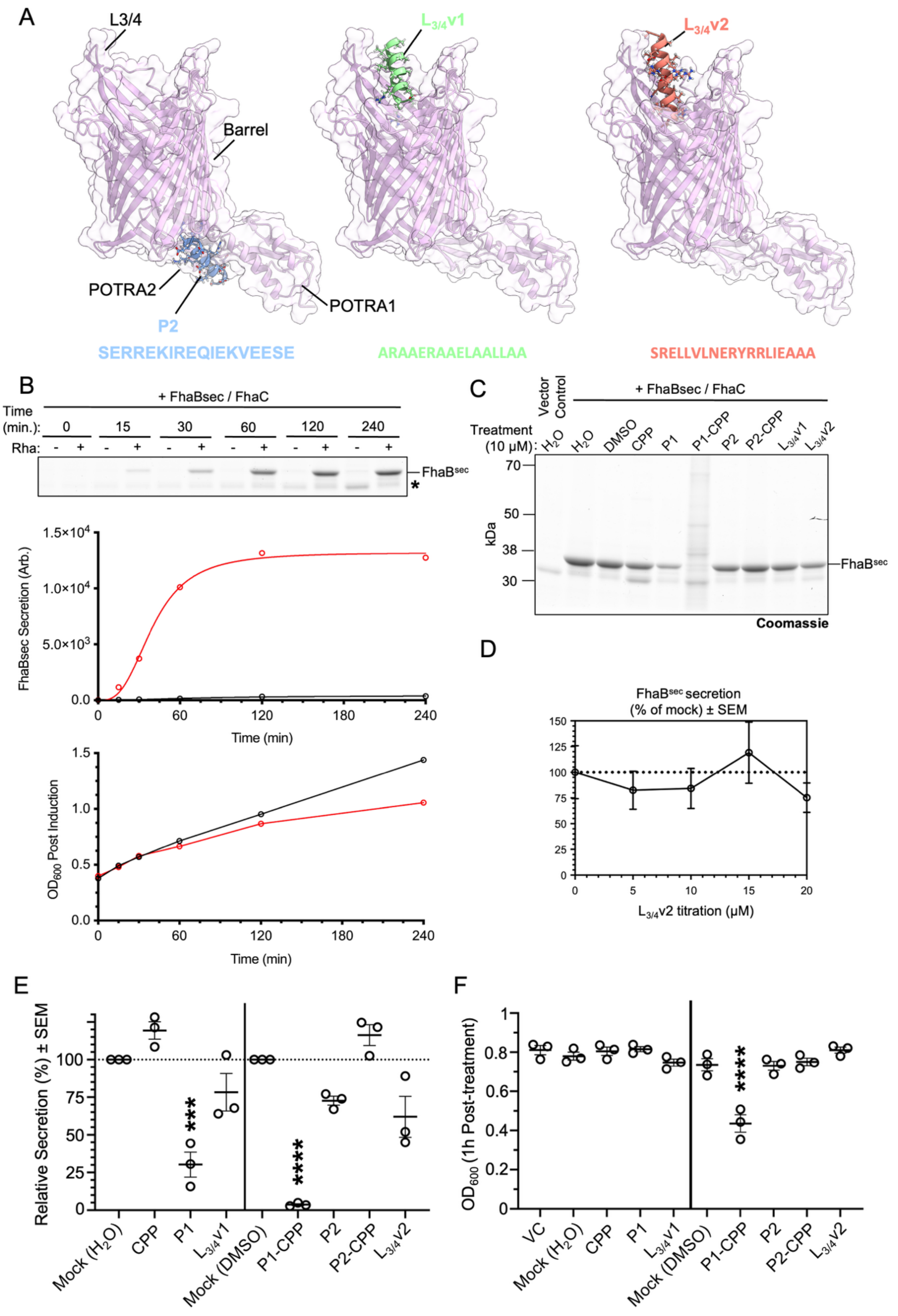
A
L3/4
Barrel
POTRA2
POTRA1
P2
SERREKIREQIEKVEESE
L3/4v1
ARAAERAAELAALLAA
L3/4v2
SRELLVLNERYRRLIEAAA
B
+ FhaBsec / FhaC
Time (min.):
0
15
30
60
120
240
Rha:
FhaBsec
*
FhaBsec Secretion (Arb.)
Time (min)
OD600 Post Induction
Time (min)
C
Vector Control
+ FhaBsec / FhaC
Treatment (10 μM):
H2O
DMSO
CPP
P1
P1-CPP
P2
P2-CPP
L3/4v1
L3/4v2
kDa
70
50
38
30
FhaBsec
Coomassie
D
FhaBsec secretion (% of mock) ± SEM
L3/4v2 titration (μM)
E
Relative Secretion (%) ± SEM
Mock (H2O)
CPP
P1
L3/4v1
Mock (DMSO)
P1-CPP
P2
P2-CPP
L3/4v2
F
OD600 (1h Post-treatment)
VC
Mock (H2O)
CPP
P1
L3/4v1
Mock (DMSO)
P1-CPP
P2
P2-CPP
L3/4v2

**Figure S2: Activity of RFdiffusion-predicted peptide binders of FhaC.**

**(A)** Predicted binding sites and sequences of other RFdiffusion derived peptides: (Left) **P2** is predicted to bind to the POTRA2 domain. (Middle) **$L_{3/4}$v1** and (Right) **$L_{3/4}$v2** are predicted to bind to extracellular loops 3 and 4. L3/4 = Extracellular Loop 3 and 4, H1 = Helix-1. **(B)** Coomassie gel of protein precipitates of culture media supernatant at each time point. An unknown endogenously released *E. coli* protein is noted by (*). $FhaB^{sec}$ was quantified and values of $FhaB^{sec}$ levels and $OD_{600}$ are plotted below for each time point. **(C)** Assay was conducted as in Figure 1C,D but with peptides added directly before rhamnose. Representative Coomassie gel from n = 3. An unknown endogenously released *E. coli* protein is noted by (*). **(D)** Secreted $FhaB^{sec}$ in **C** quantified as a mean percentage of mock treatment group. **(E)** Experiment as in **C**. Quantitation of $FhaB^{sec}$ secreted from cultures treated with different concentrations of **$L_{3/4}$v2**. Symbols represent mean percentage of mock treatment group (n = 3). **(F)** $OD_{600}$ of subcultures 1 hour after adding peptides and L-rhamnose. Line denotes the mean $OD_{600}$ value. Ordinary one-way ANOVA (with Dunnett's multiple comparisons post-hoc tests) was conducted, ****$P < 0.0001$. ANOVA statistics in Table S8.

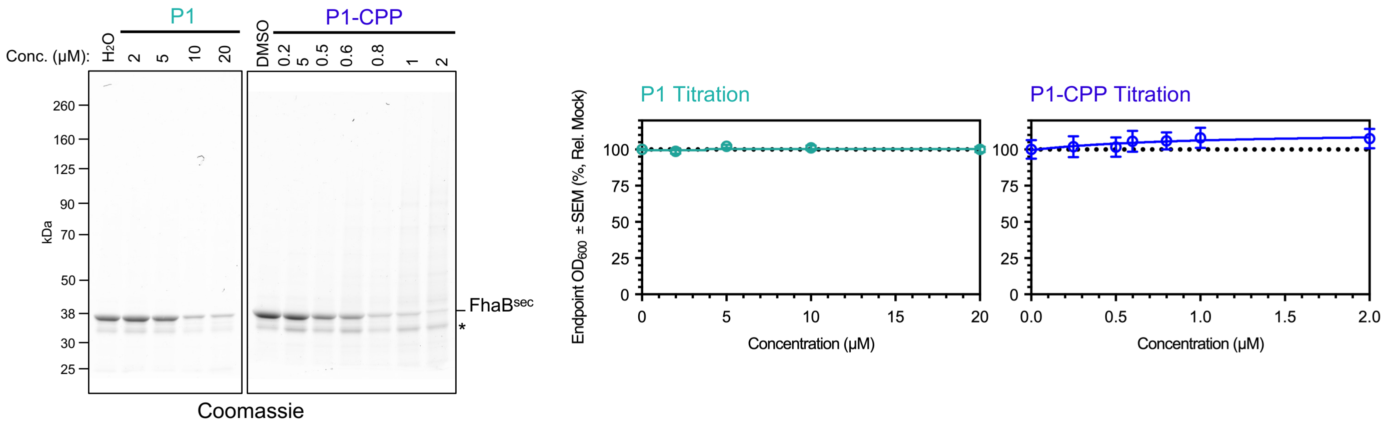


**Figure S3: Concentration-dependent inhibition of FhaC – full gels & $OD_{600}$ per concentration.**

(Left) Representative Coomassie-stained gels of samples treated with titrated **P1** and **P1$^{CPP}$** (n = 3). Assay was conducted as in **Figure 1D**. Conc. = Concentration. An unknown endogenously released *E. coli* protein is noted by (*). (Right) $OD_{600}$ of subcultures 1 hour after adding peptides at different concentrations (n = 3). Plots denote the mean $OD_{600}$ value.

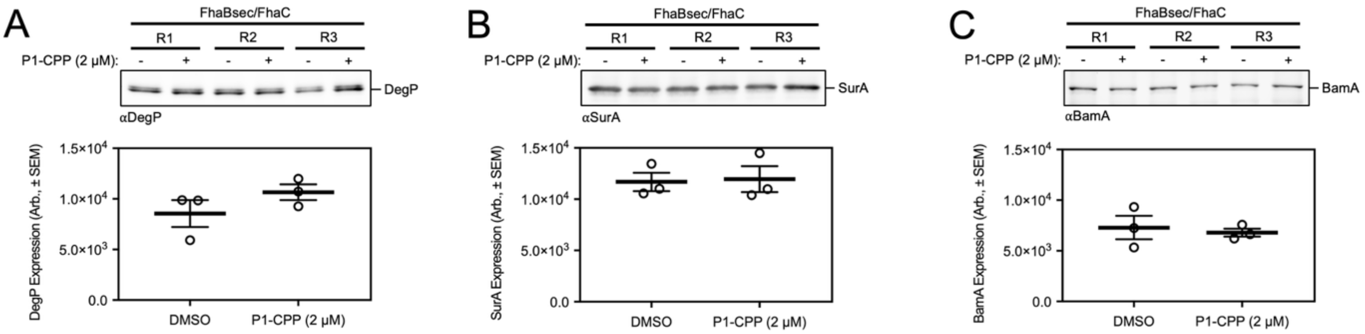


**Figure S4: P1$^{CPP}$ does not alter expression of SurA, DegP, or BamA.**

Experiments conducted as in Figure 2C. Proteins were separated by SDS-PAGE and probed by immunoblotting with αDegP (**A**), αSurA (**B**), or αBamA (**C**).

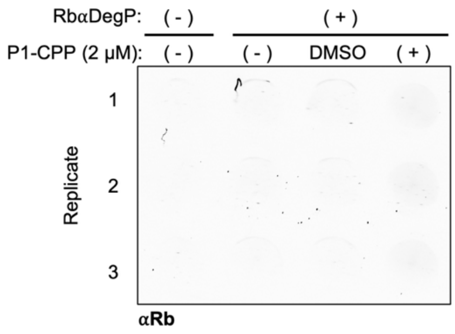


**Figure S5: Live-cell helix-mobility assay specificity control.**

Experiment conducted as in Figure 3A,B except that bacteria were probed with αDegP and an appropriate secondary antibody. Lack of detection of periplasmic DegP strongly suggests the requirement for antigens to be surface exposed for detection.

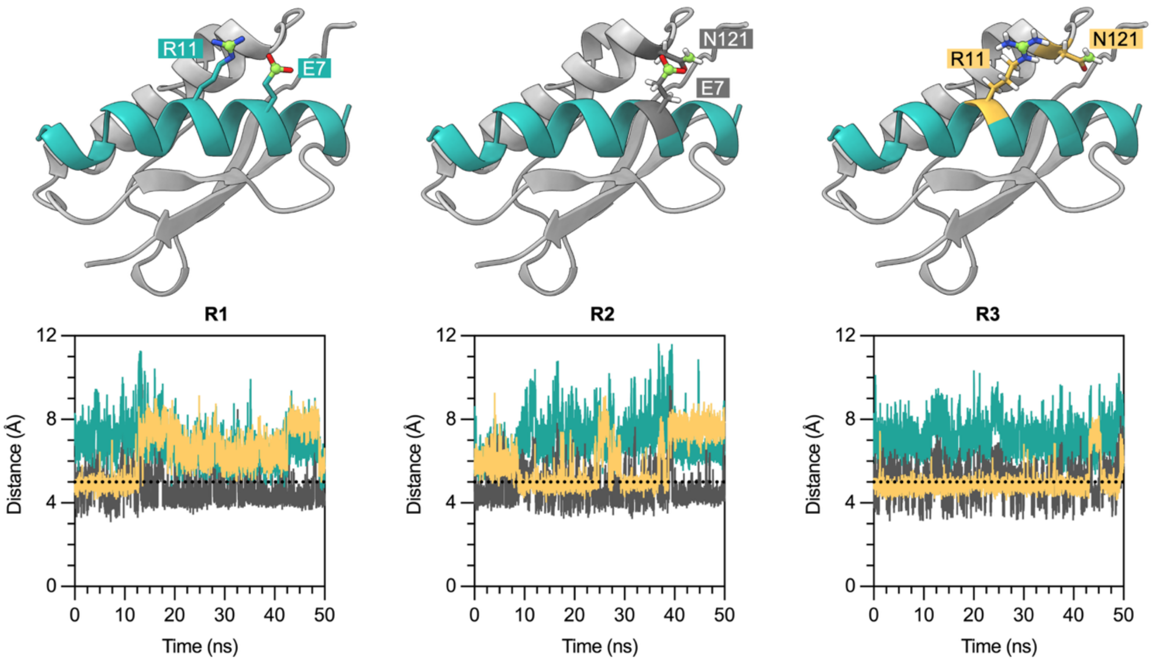


**Figure S6: P1 residue R11 interacts interchangeably with FhaC N121.**

Top row, different interactions between P1(E7) P1(R11) and FhaC(N121) identified during MD analysis. Bottom row, Replicate all-atom MD simulations as in Figure 5B. Distance between terminal residue carbons of residue pairs as depicted as green spheres in **A**.

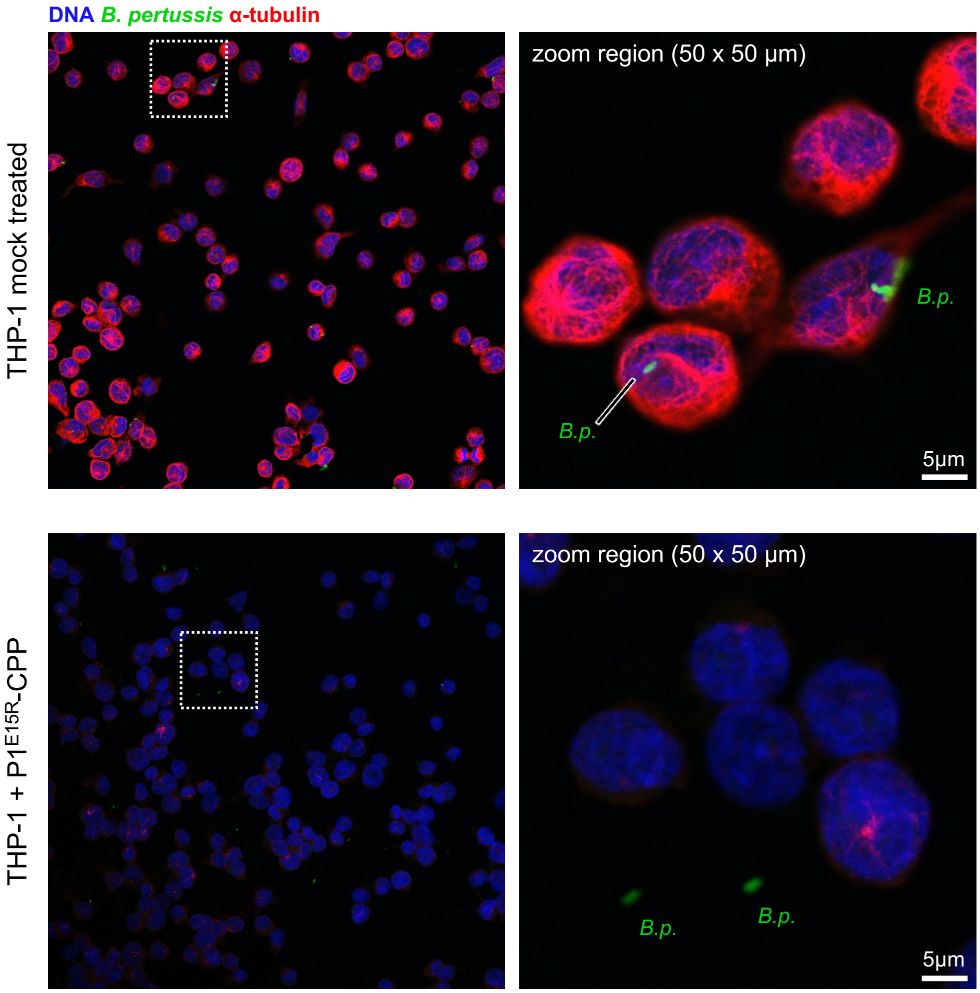


**Figure S7: P1$^{E15R\text{-}CPP}$ prevents *B. pertussis* induced cytotoxicity and adherence to differentiated macrophages.**
Experiment as in Figure 6F-G except macrophage differentiated THP-1 cell line was infected with *B. pertussis* L1423 in the presence of inhibitor treatment (or mock treatment) and subjected to fluorescence microscopy. Representative micrographs of n = 3.

**Table S1. RFdiffusion output scores and predicted properties of the top 10 peptide sequences of each run conducted.**

Abbreviations: pLDDT = predicted local distance difference test, i_pTM = binding interface predicted template modelling, i_pAE = binding interface predicted alignment of error; RMSD = root mean square deviation, pI = Isoelectric point, GRAVY = grand average of hydropathy, cLogP = calculated LogP, cLogS = calculated LogS. The properties of the peptides were calculated using ProtParam (*3*), ALOGPS (*4*), and PepCalc (*5*).

| **POTRA1** | | | | | | | | | | | |
|---|---|---|---|---|---|---|---|---|---|---|---|
| No | pLDDT | i_pTM | i_pAE | RMSD (Å) | Sequence | M.W | pI | GRAVY | cLogP | cLogS | |
| 1 | 0.818 | 0.703 | 6.25 | 1.56 | VAAMAAEASARARAEFAAR | 1920.18 | 9.48 | 0.295 | -1.36 | -4.72 | **P1** |
| 2 | 0.753 | 0.600 | 7.58 | 2.08 | IAEMAREASARARAEFAAR | 2077.35 | 9.45 | -0.300 | -1.39 | -4.84 | |
| 3 | 0.730 | 0.534 | 8.13 | 1.38 | VAAMLAEASARARAEFAAR | 1962.26 | 9.48 | 0.400 | -1.26 | -4.84 | |
| 4 | 0.737 | 0.540 | 8.26 | 1.00 | VAEMAAEASARAREWLAAQ | 2031.27 | 4.79 | -0.021 | -1.15 | -4.89 | |
| 5 | 0.718 | 0.525 | 8.30 | 1.50 | VAEMAREASARARAELAAR | 2029.3 | 9.42 | -0.263 | -1.57 | -4.7 | |
| 6 | 0.723 | 0.544 | 8.45 | 1.34 | VAAMAAEASARARAAFAAE | 1835.07 | 6.11 | 0.626 | -1.28 | -4.81 | |
| 7 | 0.686 | 0.483 | 8.61 | 1.96 | VAELLREASERARAEFAAR | 2145.41 | 6.26 | -0.389 | -1.27 | -4.86 | |
| 8 | 0.697 | 0.476 | 8.62 | 2.41 | IAELLAEASARAREEFAAR | 2074.32 | 4.95 | -0.042 | -1.18 | -4.84 | |
| 9 | 0.652 | 0.510 | 8.70 | 1.00 | VAEMLREASERAREEFERE | 2337.55 | 4.6 | -1.33 | -1.40 | -4.54 | |
| **POTRA2** | | | | | | | | | | | |
| No | pLDDT | i_pTM | i_pAE | RMSD (Å) | Sequence | M.W | pI | GRAVY | cLogP | cLogS | |
| 1 | 0.781 | 0.645 | 7.14 | 4.69 | SERREKIREQIEKVEESE | 2274.47 | 4.86 | -2.09 | -2.14 | -4.09 | **P2** |
| 2 | 0.761 | 0.608 | 7.76 | 4.83 | SERREQIRKQIEEVEESE | 2274.43 | 4.59 | -2.07 | -1.94 | -4.03 | |
| 3 | 0.741 | 0.598 | 7.88 | 4.46 | SERREEIRRQIEKVEEEE | 2344.52 | 4.69 | -2.28 | -1.81 | -4.01 | |
| 4 | 0.720 | 0.509 | 8.95 | 4.21 | SMRRLEIEEQIKKVEESA | 2175.49 | 5.06 | -1.04 | -1.91 | -4.43 | |
| 5 | 0.706 | 0.517 | 9.03 | 4.86 | SAAREEIRKQIEAVKASE | 2015.25 | 6.02 | -0.861 | -2.22 | -4.36 | |
| 6 | 0.688 | 0.510 | 9.32 | 4.93 | SAAREEIRKQIEAVKESE | 2073.29 | 5.06 | -1.16 | -2.08 | -4.28 | |
| 7 | 0.642 | 0.494 | 9.67 | 4.67 | EERREEIRKQIEEVEASE | 2259.41 | 4.47 | -1.93 | -1.79 | -4.19 | |
| 8 | 0.653 | 0.502 | 9.74 | 3.51 | EERRKEIEEQIKKVEESE | 2288.5 | 4.69 | -2.21 | -2.09 | -4.3 | |
| 9 | 0.650 | 0.494 | 9.82 | 4.10 | SERREEIEKQIKEVEESE | 2247.4 | 4.47 | -2.04 | -2.06 | -4.12 | |
| 10 | 0.665 | 0.477 | 9.97 | 5.16 | SALREEIRKQIEAVEASE | 2058.28 | 4.65 | -0.728 | -1.81 | -4.34 | |
| **Extracellular Loops 3 and 4 (Try 1)** | | | | | | | | | | | |
| No | pLDDT | i_pTM | i_pAE | RMSD (Å) | Sequence | M.W | pI | GRAVY | cLogP* | cLogS* | |
| 1 | 0.374 | 0.396 | 19.4 | 41.5 | ARAAERAAELAALLAA | 1567.81 | 6.19 | 0.725 | -1.36 | -4.81 | **$L_{3/4}$v1** |
| 2 | 0.241 | 0.318 | 20.9 | 41.7 | AAAAARAAAAAAAAAA | 1240.38 | 9.79 | 1.41 | -1.91 | -4.58 | |
| 3 | 0.334 | 0.330 | 21.0 | 41.0 | ARAAARAAELAALLAA | 1509.77 | 9.64 | 1.06 | -1.38 | -4.81 | |
| 4 | 0.251 | 0.282 | 22.7 | 40.8 | AAAAARAAEAAALAAA | 1340.50 | 6.05 | 1.20 | -1.56 | -4.80 | |
| 5 | 0.251 | 0.282 | 22.7 | 40.8 | AAAAARAAEAAALAAA | 1340.50 | 6.05 | 1.20 | -1.56 | -4.80 | |

| | | | | | | | | | | |
|---|---|---|---|---|---|---|---|---|---|---|
| 6 | 0.221 | 0.237 | 23.6 | 43.5 | AAAAAAAAAEA AALAAA | 1255.39 | 4.00 | 1.59 | -1.51 | -4.66 |
| 7 | 0.270 | 0.181 | 24.9 | 43.1 | AAAAERAAEL AALLAA | 1482.70 | 4.53 | 1.12 | -1.22 | -4.92 |
| 8 | 0.273 | 0.148 | 25.8 | 59.6 | AAAAARAAEL AALLAA | 1424.66 | 6.05 | 1.45 | -1.17 | -4.94 |
| **Extracellular Loops 3 and 4 (Try 2)** | | | | | | | | | | |
| No | pLDDT | i_pTM | i_pAE | RMSD (Å) | Sequence | M.W | pI | GRAVY | cLogP* | cLogS* |
| 1 | 0.781 | 0.809 | 7.40 | 0.413 | SRELLVLNERY RRLIEAAA | 2272.64 | 8.46 | -0.253 | -1.14 | -4.84 **$L_{3/4}$v2** |
| 2 | 0.762 | 0.793 | 7.98 | 0.509 | SRELLELNEQY RQLIEAAA | 2246.51 | 4.49 | -0.553 | -1.30 | -4.67 |
| 3 | 0.753 | 0.794 | 8.13 | 0.597 | SRDILELNKQFR ERIEAAA | 2259.55 | 5.98 | -0.758 | -1.63 | -4.66 |
| 4 | 0.729 | 0.787 | 8.85 | 0.616 | SFLIRELNRQYR ALIEAAA | 2234.59 | 8.46 | 0.042 | -1.15 | -4.94 |
| 5 | 0.697 | 0.752 | 9.62 | 0.714 | SVDLLQLNEQF RAMIEAAA | 2119.42 | 4.14 | 0.3 | -1.18 | -4.94 |
| 6 | 0.676 | 0.732 | 10.1 | 0.745 | SRDLLALNERF RALIEAAA | 2129.45 | 5.9 | 0.168 | -1.19 | -4.93 |
| 7 | 0.585 | 0.673 | 12.1 | 2.35 | SFDLLVLNEEFK KLIEEAA | 2208.54 | 4.25 | 0.184 | -1.82 | -5.12 |
| 8 | 0.546 | 0.621 | 13.3 | 1.01 | SYDLLVLNEEF KARIEAAA | 2152.43 | 4.41 | 0.111 | -1.59 | -4.86 |

**Table S2. Molecular properties of P1 and derivatives.**

Abbreviations: M.W = molecular weight, Da = Daltons, pI = isoelectric point, GRAVY = grand average of hydropathy, CPP = cell penetrating peptide, $(KFF)_3K$ = Sequence of CPP: KFFKFFKFFK, ΔNt = Truncation of 1st six N-terminal residues of P1, ΔCt = Truncation of last four C-terminal residues of P1.

| Name | Sequence | M.W (Da) | pI | GRAVY |
|---|---|---|---|---|
| P1 | VAAMAAEASARARAEFAAR | 1920.18 | 9.48 | 0.295 |
| P1$^{\text{CPP}}$ | $(KFF)_3$KGVAAMAAEASARARAEFAAR | 3372.98 | 11.1 | 0.213 |
| P1$^{\Delta\text{Nt-CPP}}$ | $(KFF)_3$KGEASARARAEFAAR | 2858.34 | 11.1 | -0.287 |
| P1$^{\Delta\text{Ct-CPP}}$ | $(KFF)_3$KGVAAMAAEASARARAE | 2927.46 | 10.5 | 0.173 |
| P1$^{\text{R11E-CPP}}$ | $(KFF)_3$KGVAAMAAEASA**E**ARAEFAAR | 3345.91 | 9.99 | 0.247 |
| P1$^{\text{R13E-CPP}}$ | $(KFF)_3$KGVAAMAAEASARA**E**AEFAAR | 3345.91 | 9.99 | 0.247 |
| P1$^{\text{E15R-CPP}}$ | $(KFF)_3$KGVAAMAAEASARARA**R**FAAR | 3400.06 | 12.0 | 0.180 |
| P1$^{\text{R11E,R13E-CPP}}$ | $(KFF)_3$KGVAAMAAEASA**E**A**E**AEFAAR | 3318.84 | 8.43 | 0.280 |
| P1$^{\text{R11E,E15R-CPP}}$ | $(KFF)_3$KGVAAMAAEASA**E**ARA**R**FAAR | 3372.98 | 11.1 | 0.213 |
| P1$^{\text{R13E,E15R-CPP}}$ | $(KFF)_3$KGVAAMAAEASARA**E**A**R**FAAR | 3372.98 | 11.1 | 0.213 |
| P1$^{\text{R11E,R13E,E15R-CPP}}$ | $(KFF)_3$KGVAAMAAEASA**E**A**E**A**R**FAAR | 3345.91 | 9.99 | 0.247 |

**Table S3.** Strains used in this study.

| Strain | Parent | Plasmid | Note | Marker | Source |
|---|---|---|---|---|---|
| NEB5α | - | - | *E. coli* K-12 strain NEB5α | - | NEB |
| XL10-Gold | - | - | *E. coli* K-12 strain XL10-Gold | - | Agilent |
| BL21(DE3) | - | - | *E. coli* B strain BL21(DE3) | - | NEB |
| RH03 | | | *E. coli* B strain RHO3 (Δasd thi-1 thr-1 leuB26 tonA21 lacY1 supE44 recA; integrated RP4-2 Tcr::Mu ΔaphA (λpir+) | DAP (400 μg/ml) | (*6*) |
| RB50 | - | - | *B. bronchiseptica* strain RB50 | - | (*7*) |
| RBX9F | - | - | *B. bronchiseptica* strain RBX9 with *fhaB* gene knocked out (Δ*fhaB*) | - | (*8*) |
| Δ*fhaC* | - | - | *B. bronchiseptica* strain RB50 with deletions in *fhaC* gene (Δaa18-576) | - | This Study |
| L1423 | - | - | *B. pertussis* SNP cluster I *(fim3A, prn2, ptxA1, ptxP3*) [*prn+/fha+*] | - | (*9*) |
| L1756 | - | - | *B. pertussis* SNP cluster I *(fim3A, prn2, ptxA1, ptxP3*) [*prn-/fha+*] | - | (*9*) |
| L2228 | - | - | *B. pertussis* SNP cluster I *(fim3A, prn2, ptxA1, ptxP3*) [*prn-/fha-*] | - | (*10*) |

**Table S4. Plasmids used in this study.**

Abbreviations: TS = TwinStrepII-tag, His = His x 8 or 10 tag, aa = amino acid number, $BamA_{ss}$ = BamA signal sequence, $^{E}$P1 = Codons encoding $BamA_{ss}$ - P1 peptide (VAAMAAEASARARAEFAAR), H1 = FhaC Helix 1 (31-63), H1-L = FhaC Helix 1 and linker (31-86), $Ap^{R}$ = ampicillin resistance, $Tp^{R}$ = trimethoprim resistance, SS = signal sequence, $His_{10}$ = His x 10 tag.

| Name | Construct | Notes | Marker | Reference |
|---|---|---|---|---|
| pTrc99a | | IPTG-inducible plasmid | $Ap^{R}$ | Novopro |
| pSCRhaB2 | | Rhamnose-inducible plasmid | $Tp^{R}$ | (*11*) |
| pBBR1MCS 2-fuGFP | pBBR1MCS2-fuGFP | pBBR1MCS-2 plasmid to constitutively express free-use green fluorescent protein (fuGFP) | $Km^{R}$ | (*9*) |
| pRJ64 | pTrc99a::$^{His}$FhaC | IPTG-inducible plasmid expressing an N-terminally His-tagged FhaC | $Ap^{R}$ | This Study |
| pRJ72 | pSCRhaB2::$FhaC_{WT}$ + $FhaBsec^{TS}$ | Rhamnose-inducible plasmid co-expressing wild-type FhaC and a truncated, hypersecreting form of FhaB ($FhaB^{sec}$) with a C-terminal TwinStrepII-tag | $Tp^{R}$ | This Study |
| pRJ94 | | Allelic-exchange plasmid with deletions in codons encoding aa18-576 in FhaC (with pSS4245 backbone) | $Dap^{R}$ | This Study |
| pMTDS109 | pSCRhaB2::$FhaC_{WT}$ + $FhaBsec^{TS}$ | Rhamnose-inducible vector co-expressing wild-type FhaC and a truncated, hypersecreting form of FhaB (FhaBsec) with a C-terminal TwinStrepII-tag | $Tp^{R}$ | This Study |
| pMTDS131 | pTrc99a::$^{His}$FhaC | IPTG-inducible vector expressing an N-terminally His-tagged FhaC | $Ap^{R}$ | This Study |
| pMTDS263 | pSCRhaB2::$FhaC_{WT}$ + $FhaBsec^{TS}$ + $^{E}$P1 | Rhamnose-inducible vector co-expressing wild-type FhaC, a truncated, hypersecreting form of FhaB (FhaBsec) with a C-terminal TwinStrepII-tag, and P1 peptide with an N-terminal $BamA_{ss}$ | $Tp^{R}$ | This Study |
| pMTDS323 | pSCRhaB2::$^{His}$FhaC + $FhaBsec^{TS}$ | Rhamnose-inducible vector co-expressing N-terminally His-tagged FhaC and a truncated, hypersecreting form of FhaB (FhaBsec) with a C-terminal TwinStrepII-tag | $Tp^{R}$ | This Study |
| pMTDS497 | pSCRhaB2:: $^{His}$FhaCΔH1-L + $FhaBsec^{TS}$ | Rhamnose-inducible vector co-expressing N-terminally His-tagged FhaC with Helix 1 and its linker truncated and a truncated, hypersecreting form of FhaB (FhaBsec) with a C-terminal TwinStrepII-tag | $Tp^{R}$ | This Study |
| pMTDS607 | pSCRhaB2:: $^{His}$FhaCΔH1 + $FhaBsec^{TS}$ | Rhamnose-inducible vector co-expressing N-terminally His-tagged FhaC with Helix 1 truncated and a truncated, hypersecreting form of FhaB (FhaBsec) with a C-terminal TwinStrepII-tag | $Tp^{R}$ | This Study |
| pMTDS608 | pSCRhaB2::$FhaC_{WT}$ + $FhaBsec^{TS}$ + $^{E}P1_{\Delta SS}$ | Rhamnose-inducible vector co-expressing wild-type FhaC, a truncated, hypersecreting form of FhaB (FhaBsec) with a C-terminal TwinStrepII-tag, and P1 peptide with its N-terminal $BamA_{ss}$ truncated | $Tp^{R}$ | This Study |
| pMTDS630 | pSCRhaB2:: $^{His}$FhaC | Rhamnose-inducible vector expressing an N-terminally His-tagged FhaC | $Tp^{R}$ | This Study |

**Table S5. Single-stranded oligonucleotides used in this study.**

Abbreviations: H1 = FhaC Helix 1 (31-63), H1-L = FhaC Helix 1 and linker (31-86), $Tp^R$ = trimethoprim resistance, GA = Gibson Assembly method, $^E$P1 = Codons encoding $BamA_{ss}$-P1 peptide (VAAMAAEASARARAEFAAR), SS = signal sequence, $His_{10}$ = His x 10 tag.

| **ssDNA Oligo** | **Sequence** | **Notes** |
|---|---|---|
| RJ481 | CGGATCTGTACACCTAGGACGCGTGCTGTACTGCATGATCGCCCTC | R', 3' flank Δ*fhaC*, add overhang with pSS4245 |
| RJ482 | TCAGAAACTGAGGCCGGCGTTGATGCAGCGCCCGAACAACCAG | R', 5' flank Δ*fhaC*, add overhang with 3' flank |
| RJ483 | CGGCCGCACTAGTGAGCTCGAATTCGTATTCATGCGCGGGACCAG | F', 5' flank Δ*fhaC*, add overhang with pSS4245 |
| RJ484 | GCCGGGCCTGGTTGTTCGGGCGCTGCATCAACGCCGGCCTCAG | R', 3' flank Δ*fhaC*, add overhang with 5' flank |
| RJ485 | CAAGCTGGTGCTGTCGCAG | F', Δ*fhaC* screening primer |
| RJ486 | CATCTACACCGGCATCGCC | R', Δ*fhaC* screening primer |
| mtd1 | TAATCATCCGGCTCGTATAATGTG | F', pTrc99a sequencing primer |
| mtd2 | GGCATGGGGTCAGGTGG | R', pTrc99a sequencing primer |
| mtd3 | CATCACGTTCATCTTTCCCTGG | F', pSCrhaB2 sequencing primer |
| mtd4 | CGGCGCTACGGCGTTTCAC | R', pSCrhaB2 sequencing primer |
| mtd64 | CGCCAACGTAAAGAGCAACTGC | F', FhaC sequencing primer (1) |
| mtd65 | GGAAGACCGGCAATATTACG | F', FhaC sequencing primer (2) |
| mtd66 | TTCAGTATAGCCGTCAGC | F', FhaC sequencing primer (3) |
| mtd67 | CAACAATGGTCTTACGGACG | F', FhaBsec sequencing primer (4) |
| mtd68 | CTTTGGGTGATGCAACGG | F', FhaBsec sequencing primer (5) |
| mtd149 | **TGT**TTACCGGGTGCACG | F', primer for FhaC *L34C* substitution |
| mtd150 | ATGATGATGATGGTGGTGATGG | R', primer for FhaC *L34C* substitution |
| mtd151 | **TGT**CGCACCATCCGTATGG | F', primer for FhaC *A533C* substitution |
| mtd152 | GTCTGGATGATTGGACTTCAGC | R', primer for FhaC *A533C* substitution |
| mtd191 | **CATCATCATCATCATCATCATCATCATCACCATGGCGGT**GCCCAGTTAT TACCGGGTGC | F', primer for FhaC $His_{10}$ tag insertion |
| mtd192 | CTGCGCACAAGCGGCTACAG | R', primer for FhaC $His_{10}$ tag insertion |
| mtd195 | CGCTCACAATTCTCAGTTTATGG | F', FhaC sequencing primer (between mtd65/66) |
| mtd196 | CTCCAAACCTGTGGGTGCTC | F', FhaC sequencing primer (between mtd66/67) |
| mtd227 | CATACAGTCACGGTTCATGCAGTG | F', primer for FhaCΔH1-L truncation |
| mtd288 | ACCGCCATGGTGATGATGATG | R', primer for FhaCΔH1-L truncation |
| mtd323 | CGCTGTAAAACATGTGTTTAGCC | F', *degP* sequencing primer |
| mtd324 | GCATCATTTCAGCCGTCAT | R', *degP* sequencing primer |
| mtd359 | CCAGTGCTGGGATGACATTA | F', *dsbA* sequencing primer |
| mtd360 | ACTGATACCTTCCGGGTTG | R', *dsbA* sequencing primer |
| mtd363 | CCAGATTTCTTGCGTACGCGATCCCCTAAGCCAAAGGTGG | F', primers to split pSCRhaB2 into 2 fragments (at the $Tp^R$ site – for GA) |
| mtd364 | AGGGGATCGCGTACGCAAGAAATCTGGTGCCGC | R', primers to split pSCRhaB2 into 2 fragments (at the $Tp^R$ site – for GA) |
| mtd369 | TCACCATGGCGGTCCTCCCGTTGAGCTGAACCC | F', primer for FhaCΔH1 truncation |
| mtd370 | GCTCAACGGGAGGACCGCCATGGTGATGATGATGATGATGATG | R', primer for FhaCΔH1 truncation |
| mtd371 | GTTTGATCCTATGGTCGCGGCCATGGCCGCC | F', primer for $^E$P1 ΔSS truncation |
| mtd372 | CCATGGCCGCGACCATAGGATCAAACCTCCTTCTAGAGGATCCCCG | R', primer for $^E$P1 ΔSS truncation |

**Table S6. Double-stranded oligonucleotide fragments used in this study.**

Abbreviations: $BamA_{ss}$ = BamA signal sequence, RBS = Ribosome binding site.

| dsDNA Fragments | Sequence | Notes |
|---|---|---|
| mtd185 | TTTTTTTTTT**tctaga**AGGAGGTTTGATCCT**ATG**GCAATG AAGAAACTTTTAATTGCCAGTTTATTGTTCTCGTCGGCGA CCGTCTACGGAGCAGTCGCGGCCATGGCCGCCGAGGCGTC TGCTCGTGCACGCGCCGAGTTTGCCGCCCGTTAA**aagctt** TTTTTTTTTT | $BamA_{ss}$-P1 peptide insert with RBS ($^{E}$P1) (flanked by HindIII/XbaI) |

### References


1. T. Maier, B. Clantin, F. Gruss, F. Dewitte, A.-S. Delattre, F. Jacob-Dubuisson, S. Hiller, V. Villeret, Conserved Omp85 lid-lock structure and substrate recognition in FhaC. *Nature Communications* **6**, 7452 (2015).
2. A. Ben Chorin, G. Masrati, A. Kessel, A. Narunsky, J. Sprinzak, S. Lahav, H. Ashkenazy, N. Ben-Tal, ConSurf-DB: An accessible repository for the evolutionary conservation patterns of the majority of PDB proteins. *Protein Science* **29**, 258-267 (2020).
3. M. R. Wilkins, E. Gasteiger, A. Bairoch, J. C. Sanchez, K. L. Williams, R. D. Appel, D. F. Hochstrasser, Protein identification and analysis tools in the ExPASy server. *Methods Mol Biol* **112**, 531-552 (1999).
4. I. V. Tetko, J. Gasteiger, R. Todeschini, A. Mauri, D. Livingstone, P. Ertl, V. A. Palyulin, E. V. Radchenko, N. S. Zefirov, A. S. Makarenko, V. Y. Tanchuk, V. V. Prokopenko, Virtual computational chemistry laboratory--design and description. *J Comput Aided Mol Des* **19**, 453-463 (2005).
5. S. Lear, S. L. Cobb, Pep-Calc.com: a set of web utilities for the calculation of peptide and peptoid properties and automatic mass spectral peak assignment. *J Comput Aided Mol Des* **30**, 271-277 (2016).
6. M. López Carolina, A. Rholl Drew, A. Trunck Lily, P. Schweizer Herbert, Versatile Dual-Technology System for Markerless Allele Replacement in Burkholderia pseudomallei. *Applied and Environmental Microbiology* **75**, 6496-6503 (2009).
7. P. A. Cotter, J. F. Miller, BvgAS-mediated signal transduction: analysis of phase-locked regulatory mutants of Bordetella bronchiseptica in a rabbit model. *Infection and Immunity* **62**, 3381-3390 (1994).
8. E. Mason, M. W. Henderson, E. V. Scheller, M. S. Byrd, P. A. Cotter, Evidence for phenotypic bistability resulting from transcriptional interference of bvgAS in Bordetella bronchiseptica. *Molecular Microbiology* **90**, 716-733 (2013).
9. S. Octavia, V. Sintchenko, G. L. Gilbert, A. Lawrence, A. D. Keil, G. Hogg, R. Lan, Newly Emerging Clones of Bordetella pertussis Carrying prn2 and ptxP3 Alleles Implicated in Australian Pertussis Epidemic in 2008–2010. *The Journal of Infectious Diseases* **205**, 1220-1224 (2012).
10. Z. Xu, S. Octavia, L. D. W. Luu, M. Payne, V. Timms, C. Y. Tay, A. Keil, V. Sintchenko, N. Guiso, R. Lan, Pertactin-Negative and Filamentous Hemagglutinin-Negative Bordetella pertussis, Australia, 2013–2017. *Emerging Infectious Disease journal* **25**, 1196 (2019).
11. S. T. Cardona, M. A. Valvano, An expression vector containing a rhamnose-inducible promoter provides tightly regulated gene expression in Burkholderia cenocepacia. *Plasmid* **54**, 219-228 (2005).
